# The draft genome of the endangered, relictual plant *Kingdonia uniflora* (Circaeasteraceae, Ranunculales) reveals potential mechanisms and perils of evolutionary specialization

**DOI:** 10.1101/2020.01.08.898460

**Authors:** Yanxia Sun, Tao Deng, Aidi Zhang, Michael J. Moore, Jacob B. Landis, Nan Lin, Huajie Zhang, Xu Zhang, Jinling Huang, Xiujun Zhang, Hang Sun, Hengchang Wang

## Abstract

*Kingdonia uniflora*, an alpine herb, has an extremely narrow distribution and represents a model for studying evolutionary mechanisms of species that have adapted to undisturbed environments for evolutionary long periods of time. We assembled a 1,004.7-Mb draft genome (encoding 43,301 genes) and investigated the evolutionary history of *K. uniflora*, along with mechanisms related to its endangered status. Phylogenomic analyses based on 497 single copy genes confirmed the sister relationship between *K. uniflora* and *Circaeaster agrestis*, which were estimated to have diverged around 52 Mya. Proliferation of LTR retrotransposons in *K. uniflora* is estimated to occur around 2.7 Mya, coinciding with one recent uplift of the Hengduan Mountains between the late Miocene and late Pliocene. Across 12 species of monocots, early-diverging eudicots and core eudicots, *K. uniflora* showed significant overrepresentation in gene families associated with DNA repair and underrepresentation in gene families associated with stress response. Most of the plastid *ndh* genes were found to be lost not only in the plastome but also in the nuclear genome of *K. uniflora*. During the evolutionary process, the overrepresentation of gene families involved in DNA repair could help asexual *K. uniflora* reduce the accumulation of deleterious mutations, while at the same time, reducing genetic diversity which is important in responding to environment fluctuations. The underrepresentation of gene families related to stress response and functional loss of *ndh* genes could be due to lack or loss of ability to respond to environmental changes caused by long-term adaptation to a relatively stable ecological environment.

## Introduction

Habitat destruction caused by changing climate and human activities has driven numerous plant species to endangered status (Yang et al., 2018). Recently there has been a push to establish natural reserves to reduce the chance of extinction in vulnerable species. Nevertheless, as generally recognized, there is a need to apply genomics in conservation, employing genomic analyses to preserve plant diversity (Shafer et al., 2015). Genomics provides new opportunities for a detailed understanding of genetic diversity across the genome and the process involved in generating or losing this diversity (Windig and Engelsma, 2011), which provides aid to those making crucial policy decisions for conservation (Garner et al., 2016).

Plant lineages vary widely in their geographic distributions due to numerous factors, among which one crucial factor is adaptive capacity to respond to environmental changes. Lineages maintaining a small distribution are likely to possess low adaptive capacity to respond to geological and climatic changes at large scales, as well as habitat changes at small scales. For example, some asexual lineages, generating genetically and phenotypically identical individuals, are limited in their capacity to respond quickly to environmental fluctuation due to low genetic diversity, which can lead to a status of endangerment or even extinction (Bell and Collins, 2008). In addition, species living in an equable environment for long periods might lack or lose the ability to defend against rapidly fluctuating environmental stress. Once the habitat is altered or destroyed, these species are incapable of colonizing new habitats. Shrinking habitats and low adaptive ability to new environments together lead some plants to endangered status.

*Kingdonia uniflora* Balf. f. and W.W. Sm. (Circaeasteraceae, Ranunculales), an alpine herb (diploid, 2n=18) has a very narrow distribution (Figure S1). The habitat of *K. uniflora* represents an ecological environment of primeval forest with few disturbances. Specifically, *K. uniflora* is restricted to growing in high altitudes (~2800-4000 m), cold, damp climates with deep humus, and usually under species of *Abies*. *Kingdonia uniflora* and *Circaeaster agrestis* Maxim. together constitute the early-diverging eudicot family Circaeasteraceae (Ranunculales) (APG IV, 2016). Previous estimates showed *K. uniflora* diverged from *C. agrestis* around 52 Mya (Ruiz-Sanchez et al., 2012); although no fossil record is known for *K. uniflora*, fossil fruits similar to those of *C. agrestis* have been reported from the mid-Albian of Virginia, USA (Crane et al., 1994; Drinna et al., 1994; Sun et al., 2017). Remarkably, different from all other angiosperms, *K. uniflora* and *C. agrestis* possess an unusual dichotomous venation (Figure S2) similar to that found in ferns and *Ginkgo* (Sun et al., 2017). All above indicate an ancient relictual character for *K. uniflora*. Additionally, *K. uniflora* typically reproduces asexually, relying on rhizome systems to produce new individuals. Hence, *K. uniflora* provides an ideal model to study the evolutionary mechanisms of ancient plant lineages that have an extremely narrow distribution and rely on a highly specialized habitat.

A previous study (Sun et al., 2017) found rampant loss and pseudogenization of *ndh* genes in the *Kingdonia uniflora* plastome. The possibility of the transfer of the lost plastid segments to the nuclear genome at that time could not be determined. In the present study, we provide a *de novo* genome sequence of *K. uniflora* using both Illumina and Pacbio sequencing technologies. We aim to investigate the evolutionary history of *K. uniflora* and reveal the potential mechanisms of its evolutionary specialization.

## Results

### Genome assembly and annotation

Genome size estimation using flow cytometry suggested a haploid genome size of 1150 Mb for *K. uniflora* (Figure S3), while *k*-mer statistics indicated a similar genome size of 1170 Mb, with very low heterozygosity (Figure S4, Table S1). In the present study, we generated 236 Gb of Illumina reads and 106 Gb Pacbio reads (Table S2). A total assembly of 1,004.7 Mb (representing ~86% of the estimated genome size), consisting of 2,932 scaffolds (scaffold N50 length, 2.09 Mb; longest scaffold, 11.5 Mb) was achieved (Table 1). A total of 43,301 protein-coding genes were predicted (Table 1), among which 35,953 genes (83.03%) were functionally annotated (Table S3). In addition to protein-coding genes, various noncoding RNA sequences were identified and annotated (Table S4), including 1,124 transfer RNAs, 715 ribosomal RNAs, 125 microRNAs, and 1,751 small nuclear RNAs. The completeness of gene regions assessed by BUSCO (Benchmarking Universal Single Copy Orthologs) showed that 90.6% of the green plant single-copy orthologs were complete (Table S5).

**Table 1.** Genome assembly of *Kingdonia uniflora*.

| Genome features | Contigs/Scaffolds |
| --- | --- |
| Total length, bp | 1,004,656,313 |
| Total number of contigs | 2932 |
| Longest length, bp | 11,531,354 |
| Length of N50, bp | 2,099,369 |
| Length of N90, bp | 292,588 |
| GC content, % | 38.04 % |
| No. of genes | 43,301 |

We compared the draft genome of *Kingdonia uniflora* with the well-annotated genomes of the model plant *Arabidopsis thaliana* (Brassicaceae) and the Ranunculales species *Aquilegia coerulea* (Ranunculaceae). The genome size of *K. uniflora* is much larger than that of both references. The *K. uniflora* genome showed strong synteny with the genome of *Aq. coerulea* (Figure 1), but weak synteny with that of *A. thaliana* (Figure S5), which is not surprising given their placement in the angiosperm tree of life. The gene density in *K. uniflora* is lower than that in *Aq. coerulea* and *A. thaliana*; while the density of TEs (transposable elements) in *K. uniflora* was higher than that in *Aq. coerulea* and *A. thaliana* (Figures 1 and S5). We also compared our draft genome sequence to five other draft genomes of Ranunculales taxa, representing three of the seven Ranunculales families that have reported genome sequences available (Table S6); the quality of our assembly is comparable to that of all the five species, generating the longest N50 length and relatively fewer scaffolds. Comparatively, the genome of *K. uniflora* is larger than other Ranunculales species with sequenced genomes, such as *Aq. coerulea* (293.08 Mb), *Eschscholzia californica* (489.065 Mb) and *Macleaya cordata* (377.83 Mb), the (Table S6).

**Figure 1.**
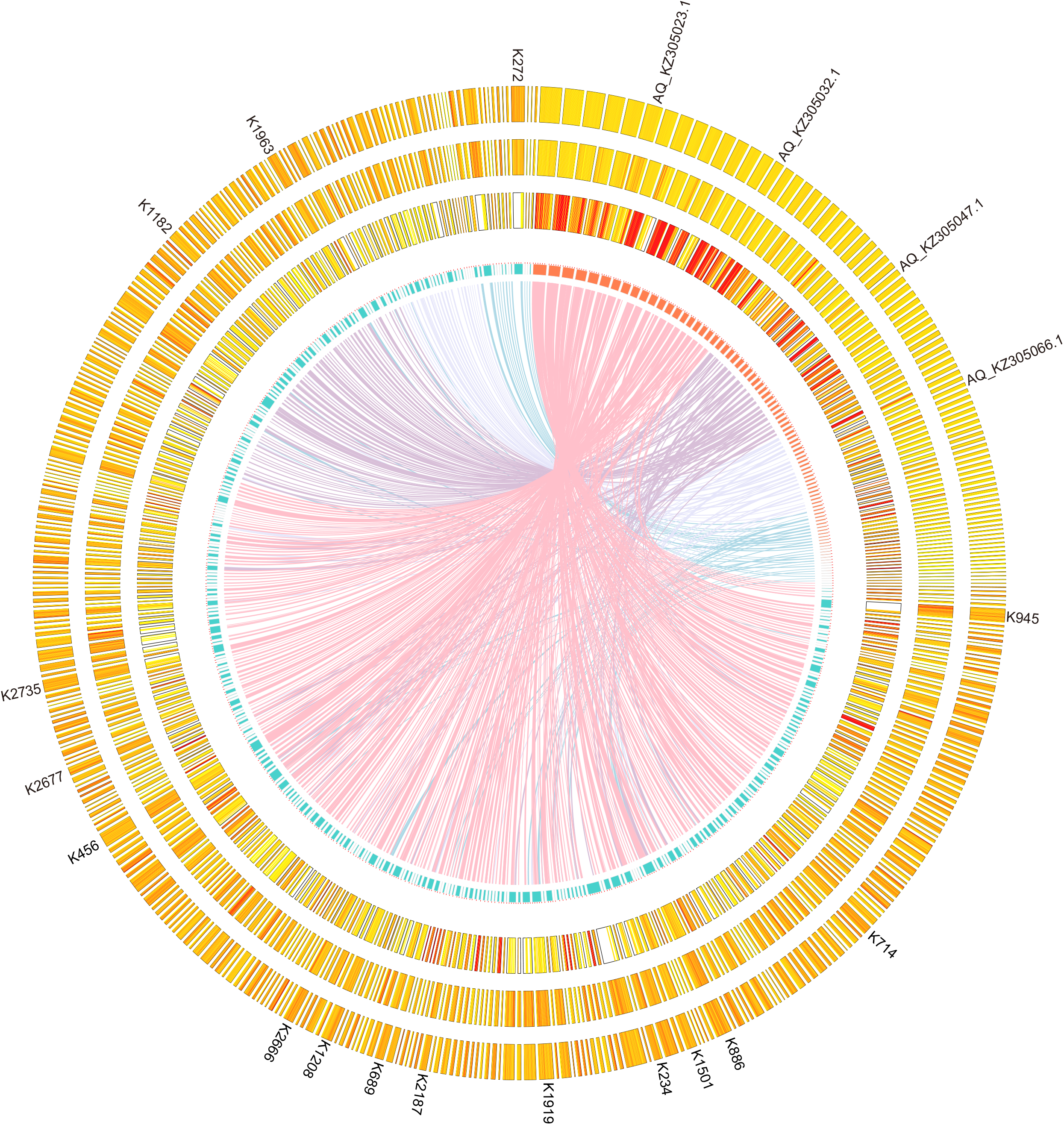
Comparative analyses of genomic features between *Kingdonia uniflora* and *Aquilegia coerulea*. Tracks from inside to outside are collinearity between both genomes, number of chromosomes/scaffolds, gene density, GC content and TE density.

### Repeat elements

Through a combination of approaches, we annotated 66.83% of the assembly as repetitive elements, among which LTRs were the most abundant, occupying 40.62% of the genome assembly length; DNA elements and long interspersed nuclear elements occupied 5.0% and 3.0% of the genome, respectively (Table S7). The proliferation of LTR retrotransposons in *K. uniflora* was estimated to peak around 2.7 Mya (Figure 2A). Analyses of age distributions built from synonymous substitutions per synonymous site (*Ks*) indicated that *K. uniflora* has undergone one recent WGD event, which occurred after its divergence from *C. agrestis* (Figure 2B). The inferred WGD event in the *K. uniflora* genome was further supported by dot-plot analysis of representative scaffolds, in which numerous paralogs derived from this event were identified (Figure 2C).

**Figure 2A.**
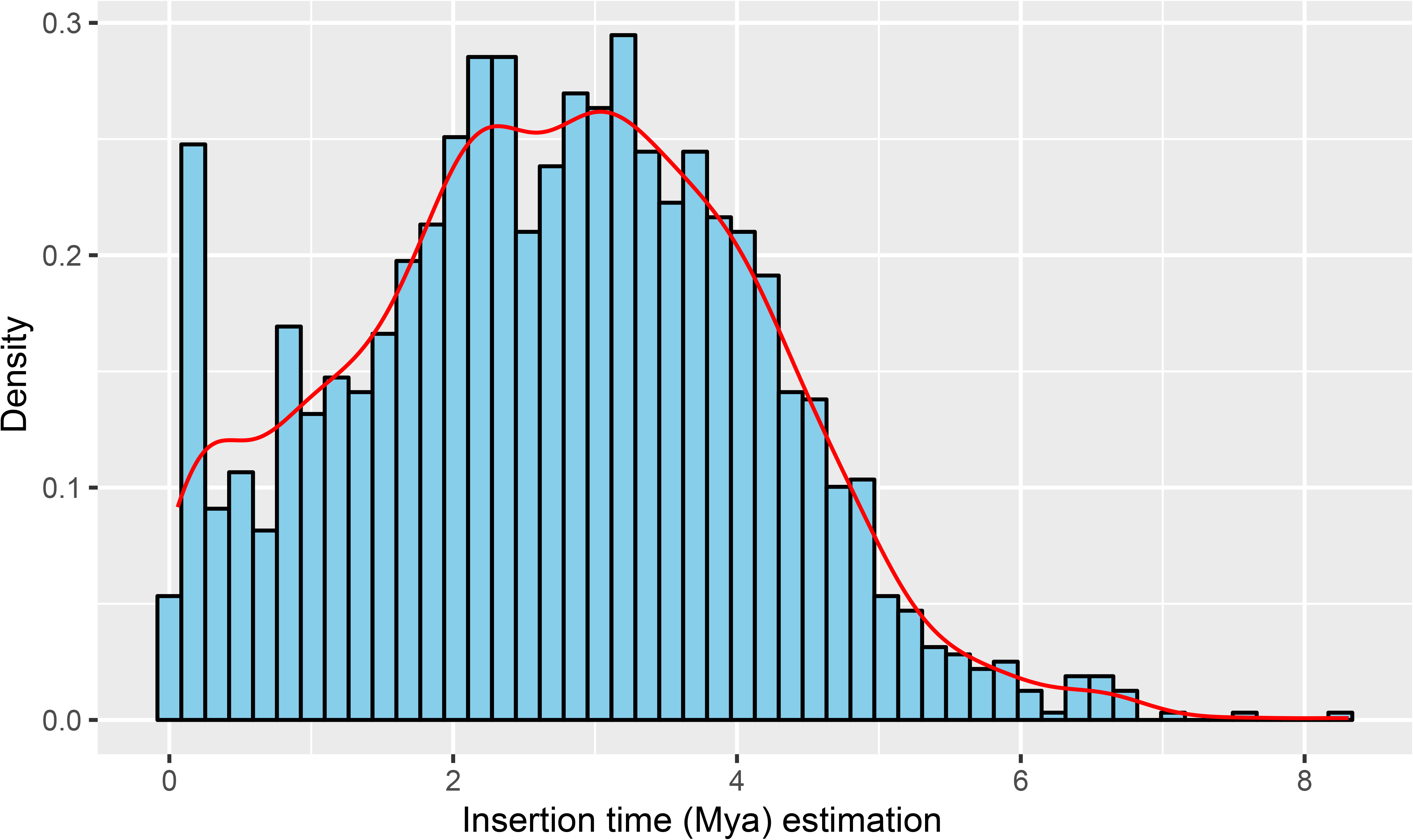
Insertion time distribution of LTR retrotransposons.

**Figure 2B.**
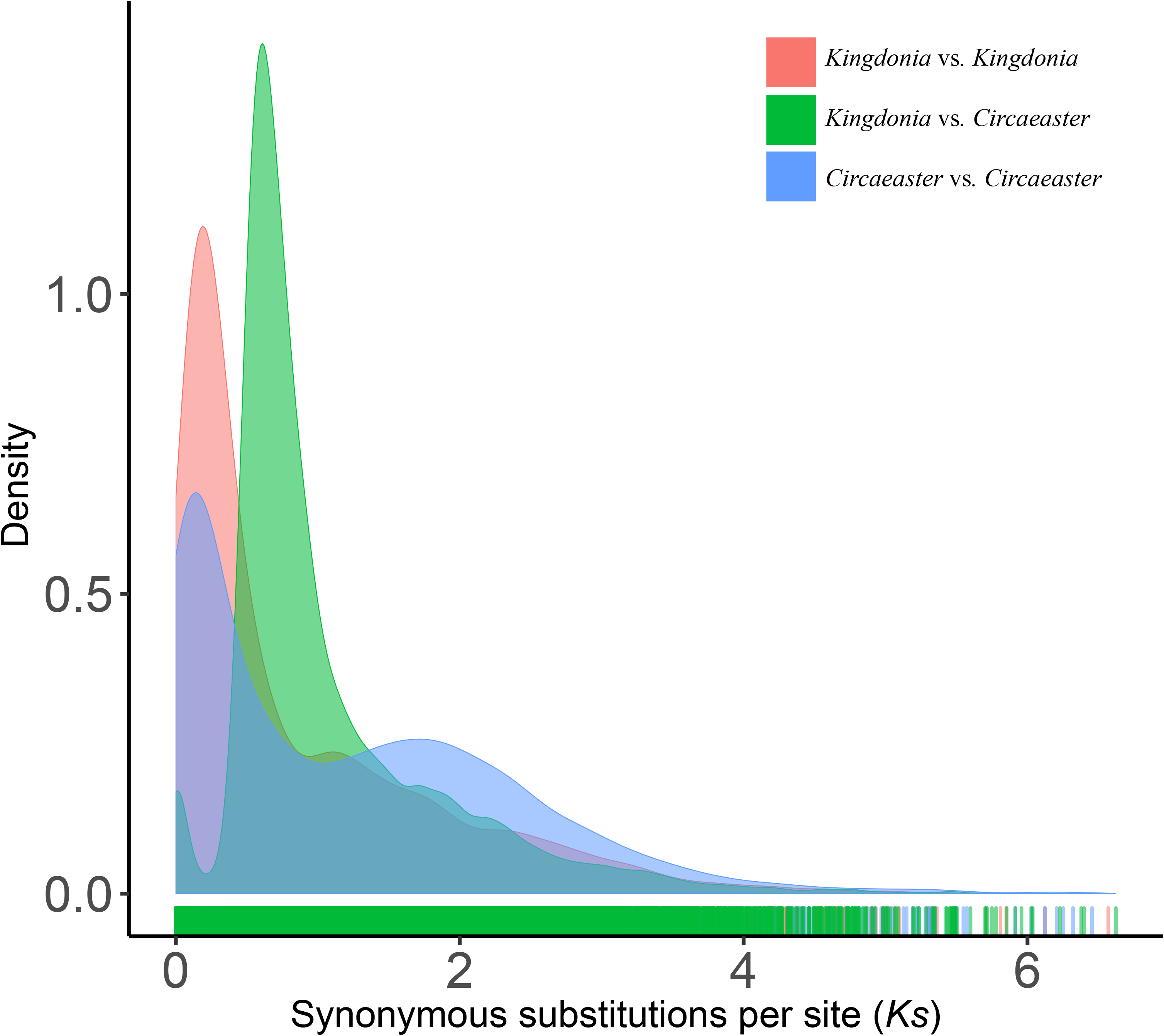
Distribution of synonymous substitution rates (*Ks*) for pairs of paralogs/orthologs in/between *K. uniflora* and *C. agrestis*.

**Figure 2C.**
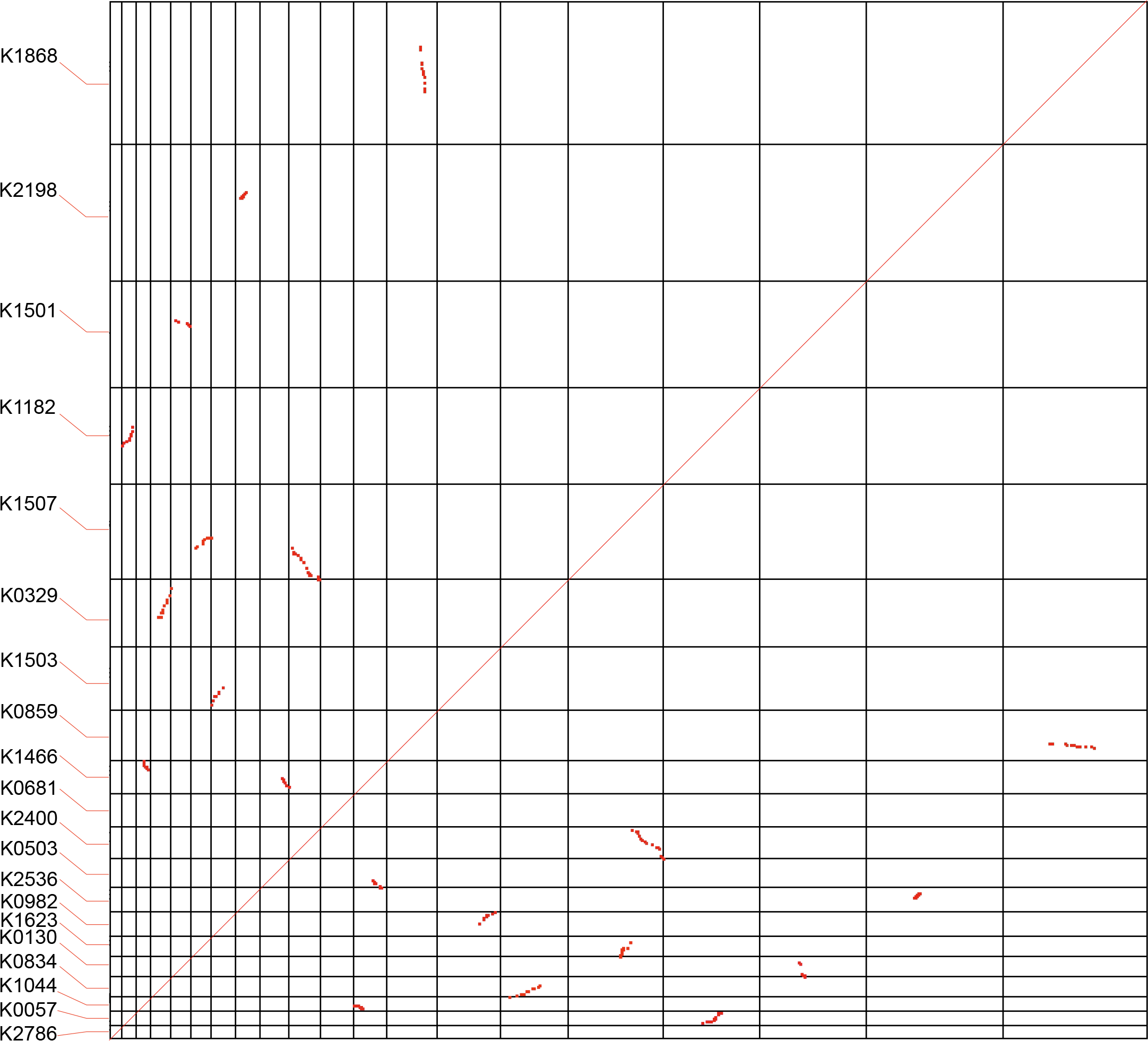
Dot plots of paralogs identified across contigs in the *K. uniflora* genome.

**Figure 2D.**
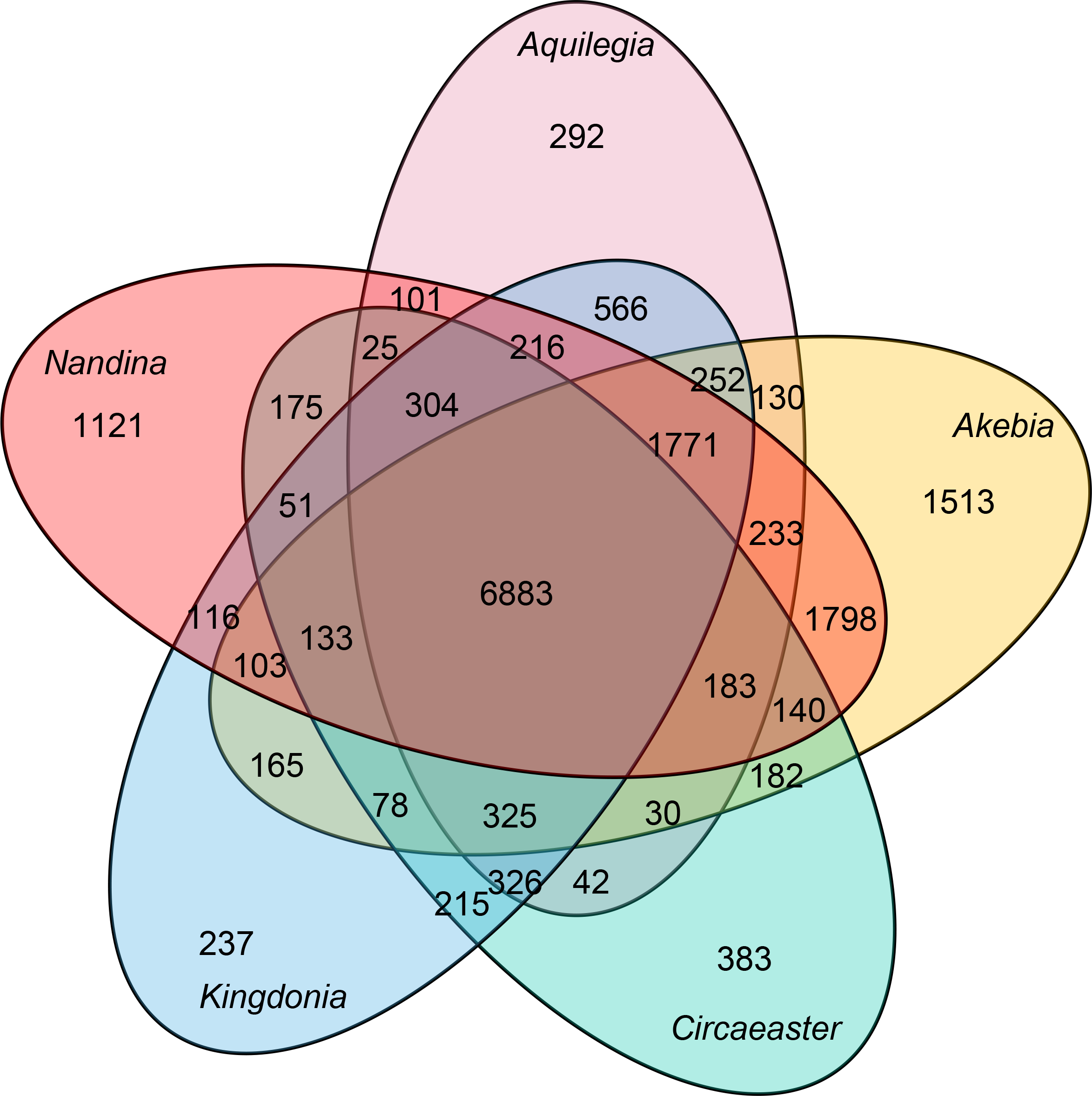
Venn diagram showing unique and shared gene families between genomes of *K. uniflora* and four other Ranunculales species.

### Phylogenetic tree construction and estimation of divergence times

Applying OrthoFinder (Emms and Kelly, 2015) to eight whole-genome and four transcriptome sequences including monocots, basal eudicots and core eudicots, we identified a total of 18,742 orthogroups, among which 6,883 were shared by *Kingdonia* and four other Ranunculales species (Figure 2D). Among these orthogroups, 497 were identified as putative single-copy gene families. To further investigate the phylogenetic relationships within Ranunculales, we conducted both concatenated and coalescent analyses using the sequences of 497 single-copy genes in 12 species. The topologies from two analyses were identical, confirming the sister relationship between *K. uniflora* and *C. agrestis*, and resolving Circaeasteraceae as sister to the clade formed by Ranunculaceae and Berberidaceae (Figure 3); Papaveraceae + Eupteleaceae was placed as the earliest-diverging clade (Figure 3). *K. uniflora* and *C. agrestis* were estimated to have diverged ~51.8 Mya in our analyses using MCMCtree with two calibration points (Figure 3). In addition, the phylogenetic analysis with an expanded group of taxa, which correspond to a larger taxonomic sampling but fewer loci indicated a similar phylogeny of Ranunculales, except the placement of Eupteleaceae (Figure S6).

**Figure 3.**
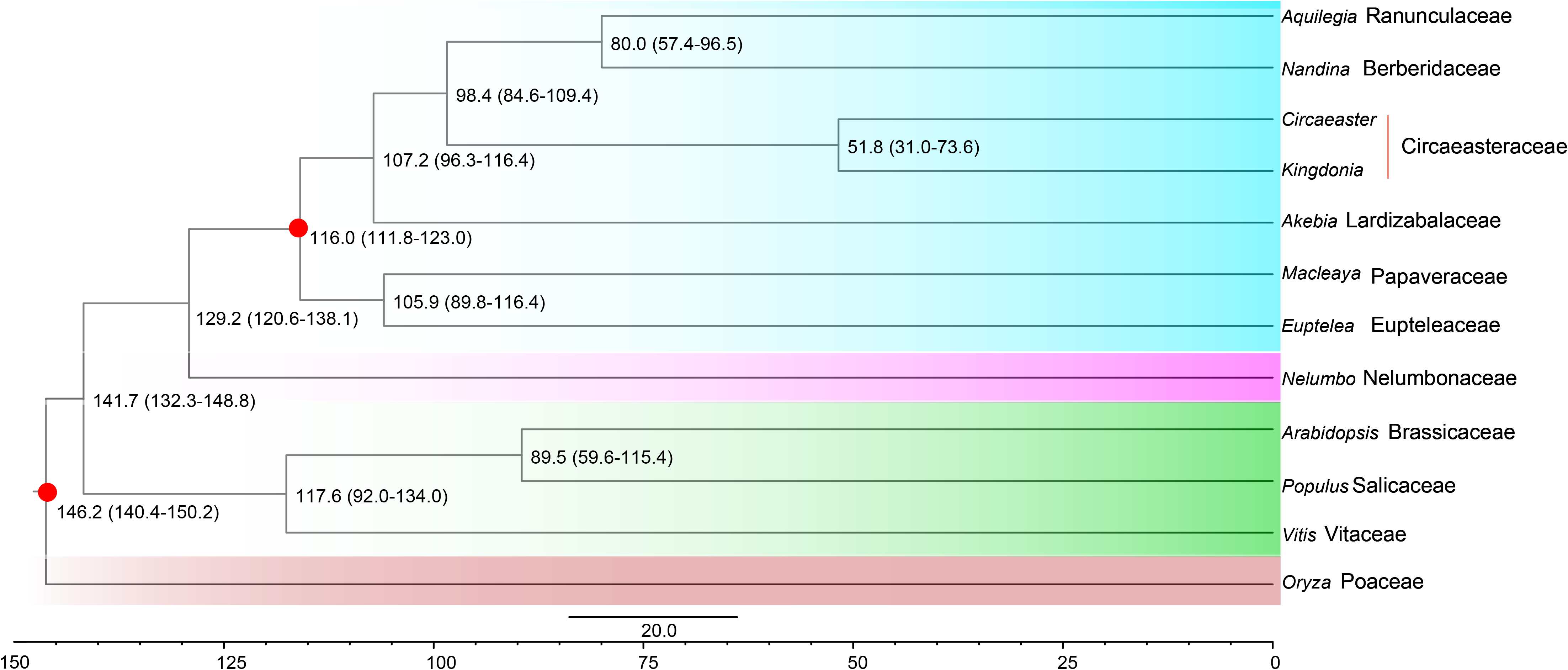
Dated phylogeny for 12 plant species with *Oryza* as an outgroup. A time scale is shown at the bottom, and red points in some nodes indicate fossil calibration points.

### Gene family overrepresentation and underrepresentation

Comparisons of the genomes among 12 species identified a total of 111 gene families that are significantly (*P*<0.01) overrepresented in *K. uniflora* and 22 gene families that are significantly underrepresented (Table S8). The results from Kyoto Encyclopedia of Genes and Genomes (KEGG) and Gene Ontology (GO) annotations showed that overrepresented gene families were considerably enriched in DNA repair pathways, such as homologous recombination, mismatch repair, DNA replication and nucleotide excision repair; while gene families showing significant underrepresentation in the *K. uniflora* genome were found to be involved in pathways related to stress or pest responses, such as the phenylpropanoid biosynthesis and secondary metabolites biosynthesis (Table 2).

**Table 2.**
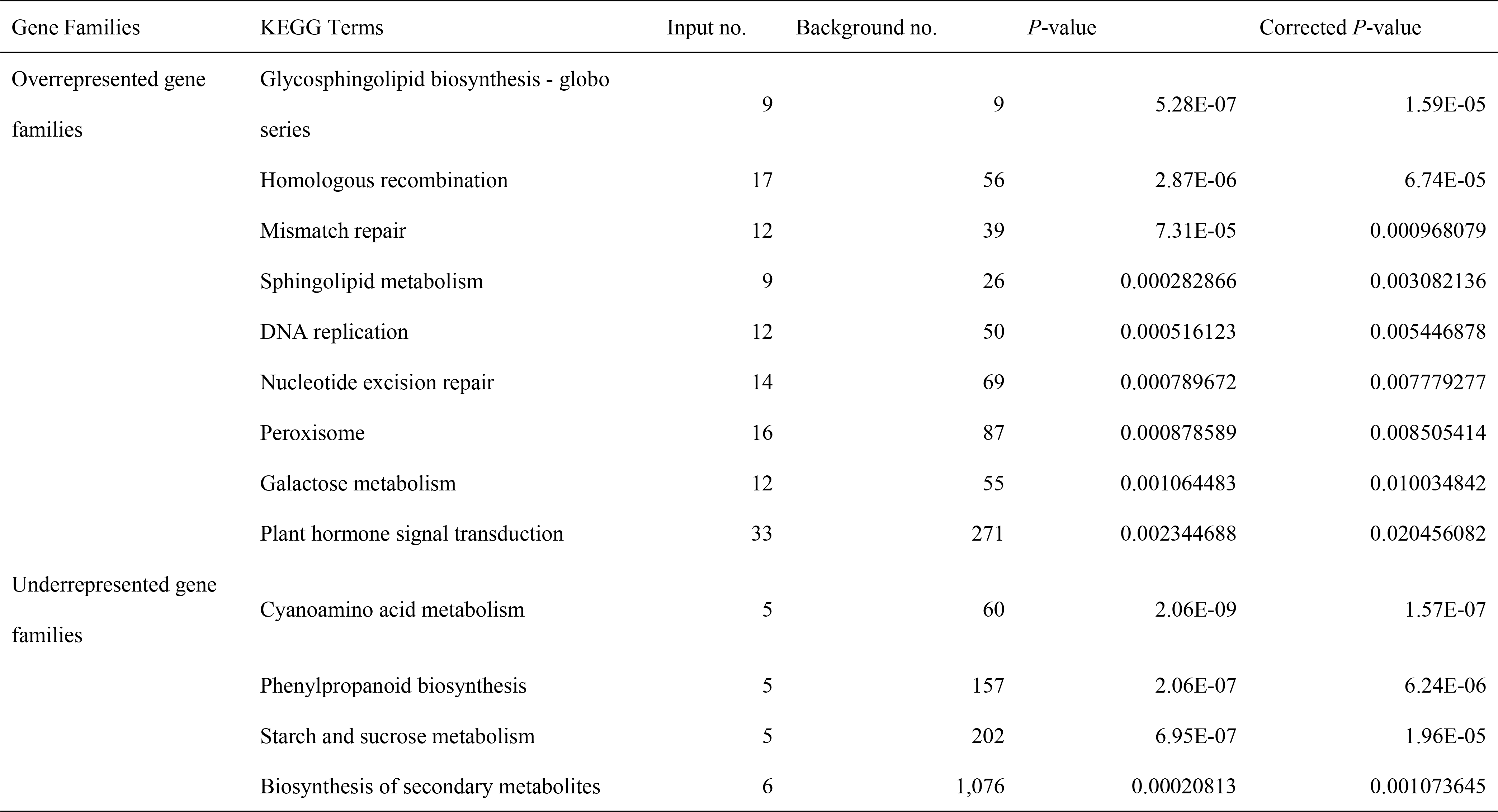
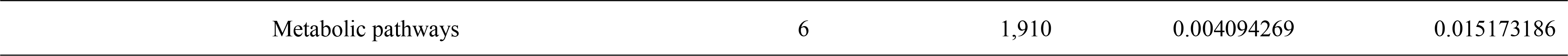
Functional annotation of the significantly overrepresented and underrepresented gene families in *Kingdonia uniflora*

### Dispensability of plastid ndh genes

To detect whether the *ndh* genes/segments lost from *K. uniflora* plastome were transferred to the nuclear genome, we conducted a BLASTN search using 11 intact *ndh* sequences extracted from *C. agrestis* as the query, using the assembled *K. uniflora* genome sequences as target. The result showed that no intact sequences for *ndh* plastid genes were discovered with the exception of *ndhE* and *ndhJ*, indicating functional copies of these genes likely have been lost (Table S9, Figure 4).

**Figure 4.**
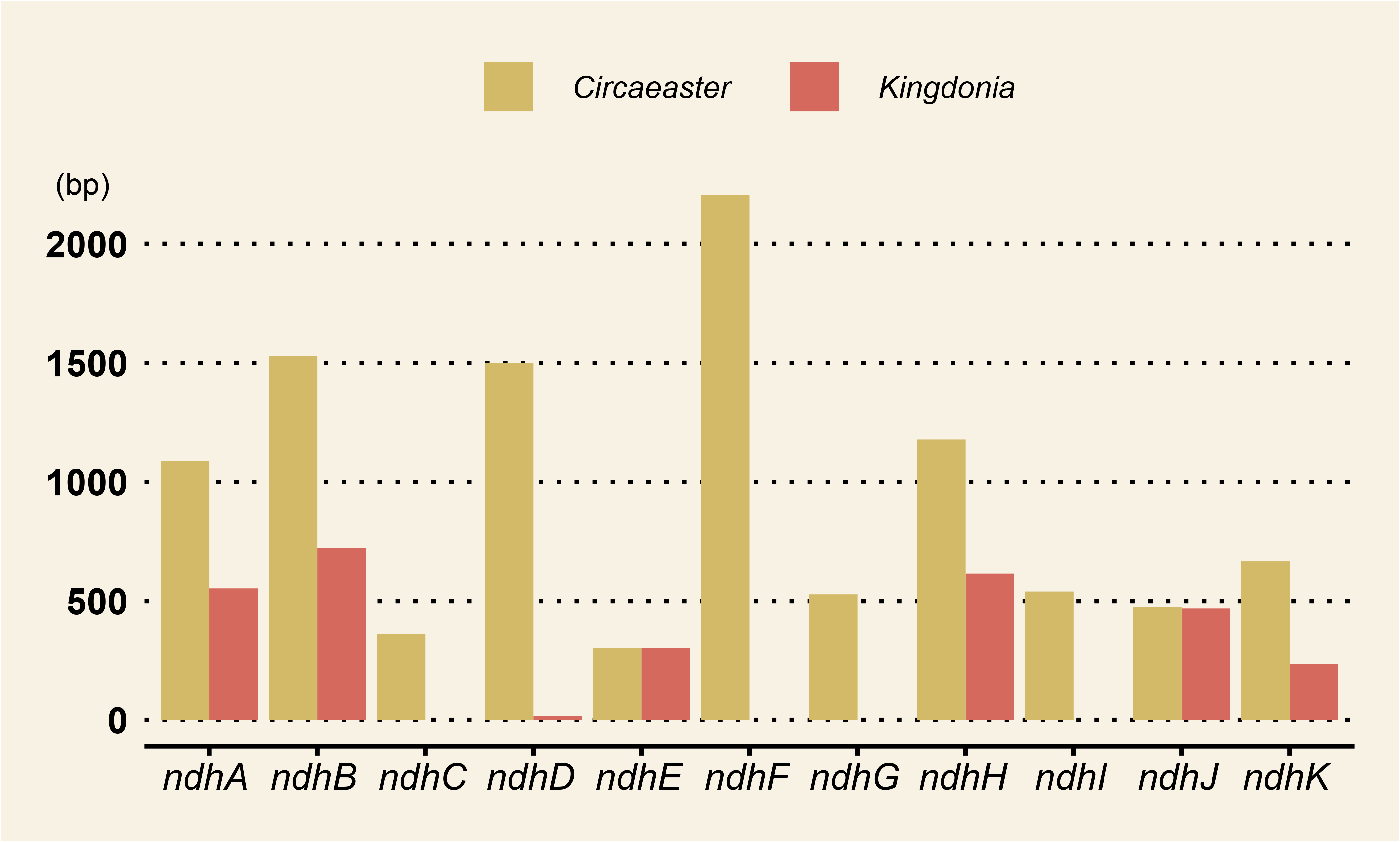
Length comparison of *ndh* genes between *K. uniflora* and *C. agrestis*.

## Discussion

Species that live in stable habitats face less stress, which can cause lack or loss of ability to respond to environmental changes. In the present study, we *de novo* assembled the genome of *K. uniflora*, an ancient relictual species exclusively found in China. Given that *K. uniflora* has a larger genome than many of its close relatives, we hypothesize that the proliferation of LTR retrotransposons and the WGD event together are likely responsible for the increased genome size of *K. uniflora* (Michael, 2014). Several studies (Tenaillon et al., 2010; Michael, 2014) have suggested the proliferation of TEs and specifically long terminal repeat retrotransposons (LTRs) in genomes is the primary driver of genome size differences in plants. This is because LTRs are expressed as RNA and reverse-transcribed into a new DNA element that can be inserted every replication cycle (Wicker et al., 2007). A comparative study using high quality genomes detected a correlation between intact LTR retrotransposons and genome size (El Baidouri et al., 2013). Abundant LTR retrotransposons were detected in the genome of *K. uniflora*, while considerably fewer LTR retrotransposons were identified in other members of the order with sequenced genomes (*Figure 1*). To investigate the evolutionary dynamics of the LTR retrotransposons, we estimated their insertion dates. The results indicate the proliferation of LTR retrotransposons in *K. uniflora* was likely triggered around one rapid uplift of the Hengduan Mountains region, occurring between the late Miocene and late Pliocene (Kirby et al., 2002; Clark et al., 2005; Sun et al., 2011; Wang et al., 2012; 2014; Meng et al., 2016; Xing and Ree, 2017). Whole genome duplications (WGD) have been shown to pervade the evolutionary history of angiosperms (Landis et al., 2018), and *K. uniflora* is no different (*Figures 2B and 2C*). Therefore, the relatively larger genome size of *K. uniflora* compared to close relatives might be promoted by both LTRs proliferation and WGD events.

Based on phylogenetic inference and estimated divergence times, we speculate that the speciation of *K. uniflora* was promoted by the Himalaya orogeny. Previous DNA based studies commonly recognized that *Kingdonia uniflora* is closely related to *Circaeaster agrestis* (Kim et al., 2004; Wang et al., 2009; Sun et al., 2017); but the relationships within Ranunculales have remained unstable (e.g., Kim et al., 2004; Wang et al., 2009; Sun et al., 2017; Lane et al., 2018). Our phylogenetic analyses confirmed the sister relationship between *K. uniflora* and *C. agrestis*, while resolving them as sister to the clade formed by Ranunculaceae and Berberidaceae, which is not congruent with previous placements of Circaeasteraceae and Lardizabalaceae as sister pairs (Kim et al., 2004; Wang et al., 2009; Sun et al., 2017). The close relationship between Ranunculaceae and Berberidaceae have been recognized in both previous and current phylogenies (e.g., Kim et al., 2004; Wang et al., 2009; Sun et al., 2017; Lane et al., 2018). However, in previous studies, either Papaveraceae (Hoot et al., 1999; Soltis et al., 2000) or Eupteleaceae (Kim et al., 2004; Wang et al., 2009; Sun et al., 2017; Lane et al., 2018) have been identified as the early-diverging clade sister to other Ranunculales lineages. Although our phylogenetic analysis based on 264 genes agree with the placement of Eupteleacea as the early-diverging clade of Ranunculales (*Figure S6*), our 497 gene set analyses resolved Papaveraceae + Eupteleaceae as the early-diverging clade (*Figure 3*). With the accumulation of molecular evidence, conflict between gene trees is ubiquitous. Conflicting gene trees could be caused by multiple factors, such as hybridization/introgression, incomplete lineage sorting, gene duplication/loss and horizontal gene transfer (Zou and Ge, 2008; Lu et al., 2016). While considering the relatively far relationships among different Ranunculales families, gene duplication/loss is possibly a major contributor to the conflicting gene trees within Ranunculales. The divergence estimation between *K. uniflora* and *C. agrestis* (~51.8 mya) is consistent with a previous estimate of ~52 Mya (Ruiz-Sanchez et al., 2012), and also coincides with the timing of the first stage of Himalayan orogeny (Rowley, 1996; Huang et al., 2015). Similar to *K. uniflora*, *C. agrestis* also has a narrow distribution confined to China and the Himalayas. However, the distribution range of *Circaeaster* is relatively larger; in places where *Kingdonia* occur *Circaeaster* can always be found; while in most regions that *Circaeaster* is distributed *Kingdonia* is absent. Previous studies have shown that orogeny could create conditions favoring speciation of resident lineages (e.g., Hoorn et al., 2013; Wen et al., 2014; Favre et al., 2015). Hence, we hypothesize that the divergence between *K. uniflora* and *C. agrestis* was likely driven by the rapid uplift of contemporary Himalayan orogeny.

Asexual reproductive system and overrepresentation of DNA repair genes together reduce genetic diversity of *K. uniflora*. Levels of genetic diversity are often associated with reproductive strategies of species (Otálora et al., 2013). Colonization, population persistence, and extinction probabilities are all influenced by the reproductive systems of species (Stephenson et al., 2000; Babará et al., 2009; Wornik and Grube, 2010; Beatty and Provan 2011; Otálora et al., 2013). *Kingdonia uniflora* primarily reproduces asexually via rhizomes. This reproductive mode could produce identical individuals rapidly, but lacks recombination and the possibility to create genetic variation in offspring, reducing the opportunities for adaptive evolution (Eckert, 2001; Castonguay and Angers, 2012). In addition, without segregation and recombination, the obligate asexual multiplication may push a species into extinction due to the steady accumulation of deleterious mutations (Thomas et al., 2016). During the long-term evolutionary history of *K. uniflora*, deleterious mutation accumulation cannot be ruled out; if something goes awry, such as the occurrence of a fatal mutation, whole clusters of clones can be wiped out. The integrated DNA-repair mechanism overrepresented in *K. uniflora* would allow it to reduce the accumulation of deleterious mutations. While, this DNA-repair system might also reduce genetic diversity produced by mutations. Having high genetic diversity is very important for plants to respond to environmental changes. We thus speculate the extremely narrow distribution range of *K. uniflora* is associated with low genetic diversity, which restricted suitable environments to simplex, equable habitats rather than multiple, divergent habitats. Specifically, *K. uniflora* can only live in high elevations with minor human disturbances, while being characterized by perennial cold temperatures of below zero degrees centigrade. The extreme temperature is likely to cause DNA, RNA, and protein damage. The overrepresentation of gene families for DNA-repair might also be one kind of protection from extremely low temperature.

Underrepresentation of genes associated with stress response in *K. uniflora* leads to degeneration of adaptive ability to environmental changes. Phenylpropanoids are believed to contribute to all aspects of plant responses towards biotic and abiotic stimuli (Vogt, 2010). As concluded by *La Camera et al.* (*2004*) when plants suffer environmental stress or pest diseases, phenylpropanoids could evoke relevant response mechanisms to protect plants from damage. Similarly, secondary metabolites also play a role in plant defense against environmental stresses and pest diseases (Bennett and Wallsgrove, 1994). Given the relatively stable ecological environment of *K. uniflora* and lack of habitat stress during growth, adaptation to such conditions resulted in the functional degeneration of stress response systems. In addition, phenylpropanoids can also promote invasion of new habitats (Bais et al., 2003; Vogt, 2010). Our results indicate the underrepresentation of gene families involved in phenylpropanoid biosynthesis, which might be one reason causing low ability of *K. uniflora* to invade new habitats.

We conclude long-term living in highly equable habitats leaded to the underrepresentation of stress response genes, which finally resulted in loss of ability to adapt to other environments; while the asexual reproductive strategy promoted overrepresentation of DNA repair genes, which reduced genetic diversity associated with adaptive capacity to environmental changes. Hence, both of the underrepresentation of stress response genes and overrepresentation of DNA repair genes are responsible for the low adaptive ability of *K. uniflora.*

Equable habitats probably promoted dispensability of most *ndh* genes in *K. uniflora*. The *ndh* genes encode subunits of the thylakoid NADPH complex that mediates cyclic electron flow around Photosystem I and facilitates chlororespiration (Martín and Sabater, 2010). Most angiosperms contain 11 plastid *ndh* genes, whereas all *ndh* genes, except for *ndhE* and *ndhJ*, were found to be either pseudogenized (*ψndhA*, *ψndhB*, *ψndhD*, *ψndhH* and *ψndhK*) or absent (*ndhC*, *ndhF*, *ndhI* and *ndhG*) in *K. uniflora* plastome (Sun et al., 2017). All 11 plastid *ndh* genes are intact in *C. agrestis* (Sun et al., 2017), indicating the loss of *ndh* genes from *K. uniflora* occurred after the split between *K. uniflora* and *C. agrestis*; suggesting that within 52 million years most of the plastid *ndh* genes were lost from *K. uniflora* not only in the plastome but also in the nuclear genome. Among land plants, the plastid *ndh* loci have also been found absent in non-photosynthetic plants, epiphytes, Gnetales, conifers and *Erodium* (Geraniaceae) (Kim et al., 2015; Lin et al., 2017; Ni et al., 2017). Evidence suggests that the thylakoid NADPH complex could optimize photosynthesis for plants under environmental stresses; while being found dispensable for plant growth under optimal growth conditions (Martín and Sabater, 2010). *Kingdonia uniflora* is extremely selective in habitat preference, and is known as the indicator for natural ecological environment without disturbance. A series of studies suggests that the *ndh* genes can be dispensable under mild non-stressing environments (e.g., Casano et al., 2001; Martín et al., 2004; Rumeau et al., 2007; Martín and Sabater, 2010). We hence speculate that the current habitats of *K. uniflora* might have promoted the dispensability of the plastid *ndh* genes. Additionally, within plants, NADPH supplies hydrogen for many anabolism processes (Antal et al., 2015). The underrepresentation of gene families related to metabolic pathways, as detected from our CAFÉ based analyses, is likely related with the nonfunction state of plastid *ndh* genes. Changing climate, shrinking habitats and low adaptive ability to environmental changes together contributed to the extremely narrow distribution of *K. uniflora.* The overrepresentation of gene families involved in DNA repair could help reduce the accumulation of deleterious mutations during asexual reproduction, which is the dominate mode of reproduction in *K. uniflora*, while at the same time, reducing genetic diversity which is important in responding to environment fluctuations. The underrepresentation of gene families in charge of stress response and nonfunction of plastid *ndh* genes are could be due to the adaptive degeneration caused by long-term adaptation to living in relatively stress free environments. Considering the long evolutionary history of *K. uniflora*, and the fossil records from *C. agrestis* in the mid-Albian of Virginia, USA (Crane et al., 1994; Drinna et al., 1994), we speculate it should have been widespread around the world. Changing climate, shrinking habitats, asexual reproduction, and adaptive degeneration caused by relying on easeful environment together lead it to a current status of endangerment.

## Methods

### Plant materials and sequencing

Fresh *K. uniflora* leaves were collected from individuals growing from the same rhizome in the Taibai Mountains (altitude 2,844 m, N 34.038°, E107.715°), Shaanxi, China. Total genomic DNA (≥10 ug, ≥50 ng/ul) was isolated from fresh leaves using the conventional cetyltriethylammonium bromide (CTAB) method (Doyle and Doyle, 1987). For Illumina sequencing, two paired-end sequencing libraries with insert sizes of 270 bp and 500 bp, respectively, were constructed and sequenced on the Illumina HiSeq X ten platform (Illumina Inc., CA, USA) at Beijing Genomics Institute (BGI) in Wuhan, Hubei, China. For PacBio single-molecule real-time sequencing, sequencing libraries with 20-kb DNA inserts were constructed and sequenced on the PacBio Sequel platform (Pacific Biosciences, CA, USA) at BGI. We also collected fresh leaves of *C. agrestis* in Taibai Mountains (altitude 2,837m, N34.038°, E107.68) for RNA extraction. Total RNA was extracted from young leaves (~100 mg) of both *K. uniflora* and *C. agrestis* using TRIzol Reagent RNA Purification (DSB, Guangdong, China). A cDNA library with insert sizes of 350-400 bp was prepared using NEBNext Ultra RNA Library Prep Kit for Illumina (NEB, MA, USA) and paired-end sequenced on the HiSeq X ten platform (Illumina Inc., CA, USA) at BGI.

### *De novo* assembly

The PacBio long reads were first corrected and *de novo* assembled using Canu v1.8 (Koren et al., 2017) with default parameters except for setting the genome size to 1.2 G to obtain contigs. Then iterative polishing was conducted on the Canu derived contigs using Pilon v1.2.3 (Walker et al., 2014) in which adapter-trimmed paired-end Illumina reads from DNA sequencing were aligned with the raw assembly with default parameters to fix bases and correct local misassembles. RNA-seq reads were assembled into transcripts using Trinity v2.6.6 (Grabherr et al., 2011) with the paired-end option and remaining default parameters.

### Annotation of repetitive sequences

We identified *de novo* repetitive sequences in the *K. uniflora* genome using RepeatModeler (http://www.repeatmasker.org/RepeatModeler/) based on a self-blast search. We further used RepeatMasker (http://www.repeatmasker.org/) to search for known repetitive sequences using a cross-match program with a Repbase-derived RepeatMasker library and the *de novo* repetitive sequences constructed by RepeatModeler. Intact LTR (long terminal repeat) retrotransposons were identified by searching the genome of *K. uniflora* with LTRharvest (Ellinghaus et al., 2008) (-motif tgca-motifmis 1) and LTR_Finder (Xu and Wang, 2007) (-D 20000 -d 1000 -L 5000-I 100). We combined results from both analyses and filtered false positives using LTR_retriever (Qu and Jiang, 2017), which also calculated the insertion date (*t*) for each LTR retrotransposons (*t*= *K/2r*, K: genetic distance) using a substitution rate (*r*) of 1.4*10−9 substitutions per site per year calculated by MCMCtree in PAML (Yang, 2007) .

### Structural and functional annotation of genes

Putative protein-coding gene structures in the *K. uniflora* genome were homology predicted using the Maker package v2.31.10 (Holt and Yandell, 2011) with protein references from the published Ranunculales genomes and the *de novo* assembled transcripts of *K. uniflora* transcriptome data generated in this study, and *de novo* predicted using Augustus v3.3.2 (Stanke et al., 2006). The rRNAs were predicted using RNAmmer v1.2 (Lagesen et al., 2007), tRNAs were predicted using tRNAscan-SE v1.4 (Lowe and Eddy, 1997), and other noncoding RNA sequences were identified using Rfam v12.0 by inner calling using Infernal v1.1.2 (Nawrocki and Eddy, 2013).

Functional annotation of the protein-coding genes was carried out by performing BLASTP analyses (e-value cut-off 1e-05) against the NCBI nonredundant protein sequence database and SwissProt. Searches for gene motifs and domains were performed using InterProScan v5.16.55 (Jones et al., 2014). Completeness of the genome was assessed by performing gene annotation using the BUSCO (v3.0.2) methods (Simão et al, 2015) by searching the Embryophyta library..

### Investigation of whole-genome duplication

We identified paralogs (within *K. uniflora* and *C. agrestis*, respectively) and orthologs (between *K. uniflora* and *C. agrestis*) using BLASTP (E value = 1E-07). For each gene pair, the number of synonymous substitutions per synonymous site (*Ks*) based on the NG method was calculated using TBtools (Chen et al., 2018); *Ks* values of all gene pairs were plotted to identify putative whole-genome duplication events. In addition, MCScanx (Wang et al., 2012) was used to identify syntenic blocks within the *K. uniflora* genome. Dot-plot analysis of syntenic blocks with at least five gene pairs was conducted using the dot plotter program within the MCScanX package to further detect whole-genome duplication events.

### Gene family and phylogenomic analysis

Orthogroups were constructed using eight genome sequences and four transcriptome sequences (Table S10). CD-HIT (Huang et al., 2010) was employed to remove redundancy caused by alternative splicing variations (-c 0.8 -aS 0.8). To exclude putative fragmented genes, genes encoding protein sequences shorter than 50 aa (amino acids) were filtered out. All filtered protein sequences of 12 species were compared with each other using BLASTP (E value = 1E-5) and clustered into orthologous groups by OrthoFinder (Emms and Kelly, 2015). Protein sequences of single-copy gene families identified by OrthoFinder were used for phylogenetic tree construction. MAFFT version 7.0 (Katoh and Standley, 2013) was used to generate multiple sequence alignment for protein sequences in each single-copy family. Poorly aligned regions were further trimmed using the Gblocks (Castresana, 2000; Talavera and Castresana, 2007). The alignments of each gene family were concatenated to a super alignment matrix, which was then used for phylogenetic tree reconstruction through the PROTCATJTT model in RAxML version 8.1.2 (Stamatakis, 2014). To assess species tree clade support, a coalescent-based analysis was also conducted using RAxML bootstrap gene trees as input for ASTRAL v. 4.7.6 (Mirarab et al., 2015). A second data set consisting of 17 taxa was also used following the same steps which consisted of increased taxonomic sampling with the tradeoff of fewer loci.

Divergence time between 12 species was estimated using MCMCtree in PAML (Yang, 2007) with the options “independent rates” and “HKY85” model. A Markov chain Monte Carlo analysis was run for 100,000,000 generations, using a burn-in of 1,000 iterations. Two constraints were used for time calibrations: (1) 140–150 Mya for the monocot-dicot split (Gaut et al., 1996; Yang et al., 2018); 112-124 Mya for the Ranunculales crown group (Magallón et al., 2015; Sun et al., 2018).

### Gene family overrepresentation and underrepresentation

Overrepresentation and underrepresentation of the OrthoFinder-derived orthologous gene families were determined using CAFÉ v. 4.1 (De Bie et al., 2006). For each significantly overrepresented and underrepresented gene family in *K. uniflora*, functional information was inferred via KOBAS (http://kobas.cbi.pku.edu.cn/anno_iden.php) using KEGG Pathway database.

### Plastid ndh gene searching

Intact **s**equences of all (11) plastid *ndh* genes, including *ndhA*, *ndhB*, *ndhC*, *ndhD*, *ndhE*, *ndhF*, *ndhG*, *ndhH*, *ndhI*, *ndhJ* and *ndhK*, were extracted from the plastome of *C. agrestis*^10^. Then BLASTN analyses (E value = 1E-5) between the 11 gene sequences and assembled *K. uniflora* genome sequences was conducted.

## Acknowledgements

This work was supported by the Strategic Priority Research Program of Chinese Academy of Sciences (XDA20050203), the Programme Foundation for the Backbone of Scientific Research by Wuhan Botanical Garden, Chinese Academy of Sciences (Y855241G01), the Major Program of National Natural Science Foundation of China (31590823), and the National Key R and D Program of China (2017YFC0505200).

