## Supplemental Tables S1-S10, Figures S1-S6 for "The draft genome of the endangered, relictual plant *Kingdonia uniflora* (Circaeasteraceae, Ranunculales) reveals potential mechanisms and perils of evolutionary specialization"

Table S1. Statistics of characteristics of *K. uniflora* genome (*K*-mer=17)

| Chracteristics | |
|---|---|
| *K*-mer | 17 |
| Peak_Depth | 80 |
| N *K*-mer | 58,788,492,58 |
| Genome size | 1170 Mb |
| Revised Genome size | 1005 Mb |
| Heterozygous rate | nan |
| Repeat rate | 0.6242 |

Table S2. Sequencing and quality filtering statistics.

| | Total data (G) | Sequence coverage (X) |
|---|---|---|
| Illumina sequencing | 236 | 201.7 |
| Pacbio Sequencing | 106.5 | 90.6 |
| Total | 342.5 | 292.7 |

Table S3. Information of function annotation in *K. uniflora* genes.

| Database | Annotated Number | Annotated Percent (%) |
|---|---|---|
| NR | 35818 | 82.7% |
| Swiss-Prot | 27859 | 64.3% |
| KEGG | 32104 | 74.1% |
| InterPro | 25951 | 59.9% |
| Total | 35953 | 83.03% |

Table S4. Statistics of noncoding RNA in *K. uniflora* genome.

| Type | | Copy | Average length (bp) | Total length (bp) |
|---|---|---|---|---|
| tRNA | | 1124 | 76 | 85487 |
| rRNA | | 715 | | |
| | 18S | 81 | 1802 | 146008 |
| | 28S | 76 | 6732 | 511650 |
| | 5.8S | 0 | _ | _ |
| | 5S | 558 | 115 | 63990 |
| snRNA | CD-box | 1447 | 104 | 149886 |
| | HACA-box | 97 | 124 | 11991 |

| | | | | |
|---|---|---|---|---|
| | splicing | 207 | 154 | 31895 |
| miRNA | | 125 | 126 | 15757 |

Table S5. Scaffolds from the *K. uniflora* assembly were aligned to conserved genes using BUSCO method.

| Species | Genome Size | BUSCO annotation assessment results |
|---|---|---|
| *K. uniflora* | 1004.7 Mb | C:90.6% [D:4.1%], F:3.3%, M:6.1%, n:1375 |

C: Complete Single-Copy BUSCOs

D: Complete Duplicated BUSCOs

F: Fragmented BUSCOs

M: Missing BUSCOs

n: Total BUSCO groups searched

Table S6. Comparison of genome assembly within Ranunculales.

| Species | Family | Genome size (Mb) | No. of scaffolds | No. of contigs | Length of N50, bp | References |
|---|---|---|---|---|---|---|
| *Aquilegia coerulea* | Ranunculaceae | 301.98 | 970 | 7,189 | 121,821 | Filiault et al., 2018 |
| *Berberis thunbergii* | Berberidaceae | 2240.74 | 11,815 | 11,815 | 397,058 | NCBI available |
| *Kingdonia uniflora* | Circaeasteraceae | 1004.7 | 2,932 | 2932 | 2,099,369 | |

| | | | | | | |
|---|---|---|---|---|---|---|
| *Eschscholzia californica* | Papaveraceae | 489.065 | 53,253 | 85,931 | 20,647 | Hori et al., 2018 |
| *Macleaya cordata* | Papaveraceae | 377.834 | 4,547 | 25,550 | 36,130 | Liu et al., 2017 |
| *Papaver somniferum* | Papaveraceae | 2715.53 | 34,381 | 65,344 | 1,773,300 | Guo et al., 2018 |

Table S7. Statistics of repeat sequences and transposable elements in *K. uniflora* genome.

| Type | Repeat size | Percent of genome (%) |
|---|---|---|
| DNA | 50812195 | 5.06 |
| LINE | 30109270 | 3.00 |
| SINE | 173746 | 0.02 |
| LTR | 408124656 | 40.62 |
| Simple repeat | 5211550 | 0.52 |
| Unknown | 179205100 | 17.84 |
| Total | 671481635 | 66.83 |

Table S8. Summary of significantly overrepresented and underrepresented orthogroups in 12 species.

| Overrepresented orthogroups | *Kingdonia* | *Arabidopsis* | *Macleaya* | *Nelumbo* | *Oryza* | *Populus* | *Vitvis* | *Circaeaster* | *Akebia* | *Aqucilegia* | *Euptelea* | *Nandina* | *P* value |
|---|---|---|---|---|---|---|---|---|---|---|---|---|---|

| | | | | | | | | | | | | | |
|---|---|---|---|---|---|---|---|---|---|---|---|---|---|
| OG0000054 | 124 | 0 | 1 | 3 | 1 | 0 | 2 | 0 | 0 | 2 | 2 | 1 | 0.0055 |
| OG0000041 | 123 | 2 | 29 | 3 | 3 | 1 | 2 | 0 | 0 | 8 | 1 | 3 | 0.0055 |
| OG0000061 | 117 | 0 | 0 | 0 | 0 | 0 | 0 | 0 | 0 | 10 | 1 | 1 | 0.004 |
| OG0000074 | 109 | 1 | 1 | 1 | 1 | 1 | 2 | 0 | 0 | 1 | 1 | 1 | 0.003 |
| OG0000098 | 102 | 0 | 0 | 0 | 0 | 0 | 0 | 0 | 0 | 0 | 0 | 0 | 0.003 |
| OG0000042 | 87 | 0 | 2 | 2 | 23 | 2 | 4 | 19 | 5 | 4 | 13 | 13 | 0.002 |
| OG0000122 | 85 | 0 | 1 | 0 | 0 | 0 | 0 | 0 | 1 | 0 | 0 | 0 | 0.002 |
| OG0000079 | 80 | 3 | 7 | 0 | 0 | 6 | 0 | 0 | 0 | 9 | 0 | 12 | 0.002 |
| OG0000023 | 79 | 22 | 14 | 12 | 14 | 28 | 11 | 1 | 19 | 50 | 2 | 5 | 0.002 |
| OG0000022 | 72 | 2 | 69 | 38 | 6 | 7 | 32 | 9 | 2 | 6 | 4 | 18 | 0.002 |
| OG0000039 | 72 | 6 | 4 | 20 | 28 | 12 | 6 | 0 | 7 | 12 | 3 | 6 | 0.002 |
| OG0000202 | 60 | 0 | 5 | 0 | 0 | 0 | 0 | 0 | 0 | 0 | 0 | 3 | 0.001 |
| OG0000247 | 60 | 0 | 0 | 0 | 0 | 0 | 0 | 0 | 0 | 2 | 0 | 0 | 0.001 |
| OG0000255 | 60 | 0 | 0 | 0 | 1 | 0 | 0 | 0 | 0 | 0 | 0 | 0 | 0.001 |
| OG0000144 | 59 | 2 | 5 | 1 | 0 | 0 | 0 | 0 | 0 | 0 | 0 | 12 | 0.001 |
| OG0000254 | 59 | 0 | 0 | 0 | 0 | 0 | 0 | 0 | 0 | 0 | 0 | 2 | 0.001 |
| OG0000096 | 58 | 4 | 2 | 3 | 3 | 6 | 2 | 4 | 5 | 10 | 4 | 3 | 0.001 |

| | | | | | | | | | | | | | |
|---|---|---|---|---|---|---|---|---|---|---|---|---|---|
| OG0000188 | 58 | 0 | 1 | 5 | 1 | 0 | 0 | 0 | 0 | 2 | 2 | 1 | 0.001 |
| OG0000317 | 53 | 0 | 0 | 0 | 0 | 0 | 0 | 0 | 0 | 2 | 0 | 0 | 0 |
| OG0000117 | 52 | 0 | 16 | 0 | 5 | 1 | 11 | 0 | 1 | 0 | 0 | 2 | 0 |
| OG0000066 | 50 | 0 | 40 | 4 | 0 | 2 | 0 | 0 | 4 | 8 | 1 | 17 | 0 |
| OG0000029 | 49 | 5 | 6 | 23 | 9 | 32 | 32 | 19 | 5 | 25 | 7 | 6 | 0 |
| OG0000290 | 48 | 1 | 1 | 0 | 0 | 0 | 0 | 0 | 1 | 4 | 1 | 1 | 0 |
| OG0000423 | 48 | 0 | 0 | 0 | 0 | 0 | 0 | 0 | 0 | 0 | 0 | 0 | 0 |
| OG0000442 | 45 | 0 | 0 | 0 | 0 | 0 | 0 | 0 | 0 | 0 | 0 | 2 | 0 |
| OG0000520 | 44 | 0 | 0 | 0 | 0 | 0 | 0 | 0 | 0 | 0 | 0 | 0 | 0 |
| OG0000223 | 41 | 3 | 0 | 0 | 13 | 0 | 0 | 0 | 0 | 7 | 0 | 0 | 0 |
| OG0000106 | 40 | 10 | 0 | 4 | 6 | 4 | 8 | 0 | 4 | 10 | 8 | 2 | 0 |
| OG0000052 | 38 | 11 | 0 | 11 | 4 | 13 | 7 | 22 | 5 | 11 | 8 | 8 | 0 |
| OG0000406 | 38 | 0 | 2 | 0 | 1 | 0 | 0 | 0 | 3 | 0 | 5 | 0 | 0 |
| OG0000610 | 35 | 1 | 2 | 0 | 0 | 1 | 1 | 0 | 0 | 1 | 0 | 0 | 0 |
| OG0000090 | 33 | 5 | 7 | 20 | 3 | 5 | 7 | 0 | 7 | 5 | 9 | 6 | 0 |
| OG0000570 | 29 | 0 | 5 | 1 | 0 | 0 | 0 | 2 | 2 | 1 | 0 | 2 | 0 |
| OG0000422 | 27 | 0 | 13 | 1 | 2 | 0 | 1 | 0 | 1 | 2 | 0 | 1 | 0 |

| | | | | | | | | | | | | | |
|---|---|---|---|---|---|---|---|---|---|---|---|---|---|
| OG0001825 | 26 | 0 | 0 | 0 | 0 | 0 | 0 | 0 | 0 | 0 | 0 | 0 | 0 |
| OG0000070 | 25 | 14 | 15 | 18 | 1 | 13 | 13 | 8 | 0 | 15 | 0 | 0 | 0 |
| OG0000439 | 24 | 3 | 2 | 0 | 6 | 1 | 0 | 0 | 1 | 3 | 2 | 5 | 0 |
| OG0000679 | 24 | 1 | 1 | 1 | 1 | 1 | 1 | 1 | 3 | 1 | 2 | 2 | 0 |
| OG0000767 | 24 | 0 | 6 | 2 | 0 | 0 | 0 | 0 | 0 | 0 | 0 | 5 | 0 |
| OG0001826 | 24 | 0 | 1 | 0 | 0 | 0 | 0 | 0 | 0 | 1 | 0 | 0 | 0 |
| OG0001538 | 23 | 0 | 0 | 0 | 0 | 0 | 0 | 0 | 2 | 2 | 0 | 1 | 0 |
| OG0002411 | 23 | 0 | 0 | 0 | 0 | 0 | 0 | 0 | 0 | 0 | 0 | 0 | 0 |
| OG0000091 | 22 | 2 | 5 | 6 | 9 | 5 | 9 | 15 | 10 | 16 | 3 | 5 | 0 |
| OG0000129 | 21 | 6 | 8 | 7 | 7 | 5 | 7 | 4 | 4 | 6 | 3 | 6 | 0.001 |
| OG0000474 | 20 | 3 | 2 | 3 | 4 | 1 | 2 | 2 | 1 | 2 | 3 | 2 | 0 |
| OG0000548 | 20 | 0 | 0 | 0 | 0 | 0 | 0 | 0 | 1 | 0 | 17 | 5 | 0 |
| OG0000705 | 20 | 3 | 2 | 1 | 1 | 2 | 3 | 0 | 4 | 1 | 0 | 1 | 0 |
| OG0000944 | 20 | 0 | 0 | 0 | 1 | 0 | 2 | 0 | 4 | 5 | 1 | 1 | 0 |
| OG0000945 | 20 | 0 | 2 | 0 | 3 | 0 | 3 | 0 | 0 | 0 | 0 | 6 | 0 |
| OG0001328 | 20 | 1 | 1 | 1 | 1 | 1 | 1 | 0 | 1 | 1 | 0 | 1 | 0 |
| OG0001010 | 19 | 1 | 1 | 1 | 1 | 1 | 1 | 2 | 1 | 1 | 2 | 2 | 0 |

| | | | | | | | | | | | | | |
|---|---|---|---|---|---|---|---|---|---|---|---|---|---|
| OG0001663 | 19 | 1 | 0 | 1 | 1 | 1 | 1 | 1 | 0 | 1 | 0 | 1 | 0 |
| OG0001823 | 19 | 0 | 4 | 0 | 0 | 0 | 0 | 2 | 0 | 0 | 1 | 0 | 0 |
| OG0002196 | 19 | 0 | 0 | 0 | 1 | 0 | 0 | 0 | 0 | 2 | 1 | 1 | 0 |
| OG0002412 | 19 | 0 | 0 | 1 | 0 | 0 | 0 | 2 | 0 | 1 | 0 | 0 | 0 |
| OG0000197 | 18 | 6 | 6 | 3 | 2 | 8 | 5 | 2 | 3 | 6 | 4 | 5 | 0.001 |
| OG0000265 | 18 | 0 | 3 | 0 | 8 | 9 | 4 | 4 | 1 | 13 | 0 | 0 | 0 |
| OG0000765 | 18 | 2 | 2 | 2 | 2 | 3 | 2 | 1 | 1 | 1 | 1 | 2 | 0 |
| OG0001389 | 18 | 1 | 1 | 1 | 1 | 2 | 1 | 0 | 0 | 1 | 1 | 2 | 0 |
| OG0003617 | 18 | 0 | 0 | 0 | 0 | 0 | 0 | 0 | 1 | 0 | 0 | 0 | 0 |
| OG0005377 | 16 | 0 | 0 | 0 | 0 | 0 | 0 | 0 | 0 | 0 | 0 | 0 | 0 |
| OG0000826 | 15 | 0 | 2 | 1 | 0 | 0 | 0 | 0 | 6 | 7 | 4 | 1 | 0 |
| OG0001084 | 15 | 1 | 1 | 1 | 2 | 1 | 1 | 1 | 2 | 1 | 4 | 2 | 0 |
| OG0001329 | 15 | 3 | 1 | 1 | 1 | 1 | 1 | 0 | 1 | 1 | 2 | 2 | 0 |
| OG0001527 | 15 | 1 | 0 | 1 | 0 | 1 | 2 | 2 | 1 | 1 | 1 | 3 | 0 |
| OG0004105 | 15 | 0 | 0 | 1 | 0 | 0 | 0 | 0 | 2 | 0 | 0 | 0 | 0 |
| OG0004682 | 15 | 0 | 0 | 0 | 0 | 0 | 0 | 1 | 1 | 0 | 0 | 0 | 0 |
| OG0006232 | 15 | 0 | 0 | 0 | 0 | 0 | 0 | 0 | 0 | 0 | 0 | 0 | 0 |

| | | | | | | | | | | | | | |
|---|---|---|---|---|---|---|---|---|---|---|---|---|---|
| OG0000182 | 14 | 12 | 1 | 4 | 7 | 6 | 6 | 6 | 2 | 10 | 2 | 2 | 0.001 |
| OG0007313 | 14 | 0 | 0 | 0 | 0 | 0 | 0 | 0 | 0 | 0 | 0 | 0 | 0 |
| OG0000206 | 13 | 10 | 6 | 5 | 3 | 9 | 5 | 0 | 3 | 6 | 2 | 5 | 0.001 |
| OG0000216 | 13 | 4 | 4 | 6 | 3 | 10 | 6 | 3 | 1 | 10 | 0 | 5 | 0 |
| OG0000543 | 13 | 1 | 1 | 6 | 4 | 4 | 1 | 1 | 4 | 7 | 0 | 1 | 0 |
| OG0001277 | 13 | 1 | 1 | 1 | 0 | 1 | 5 | 3 | 1 | 2 | 2 | 0 | 0.001 |
| OG0001985 | 13 | 2 | 1 | 1 | 1 | 1 | 2 | 0 | 1 | 0 | 2 | 1 | 0 |
| OG0003226 | 13 | 0 | 0 | 0 | 0 | 0 | 0 | 3 | 2 | 1 | 1 | 0 | 0.001 |
| OG0008474 | 13 | 0 | 0 | 0 | 0 | 0 | 0 | 0 | 0 | 0 | 0 | 0 | 0 |
| OG0000178 | 12 | 9 | 5 | 8 | 7 | 9 | 10 | 1 | 0 | 10 | 1 | 0 | 0 |
| OG0000219 | 12 | 0 | 6 | 8 | 1 | 9 | 8 | 8 | 2 | 5 | 5 | 1 | 0.001 |
| OG0000571 | 12 | 0 | 0 | 0 | 0 | 0 | 0 | 8 | 10 | 0 | 7 | 5 | 0 |
| OG0000776 | 12 | 1 | 3 | 0 | 0 | 4 | 10 | 0 | 0 | 5 | 0 | 1 | 0 |
| OG0001091 | 12 | 4 | 1 | 1 | 1 | 2 | 3 | 0 | 3 | 2 | 1 | 1 | 0.001 |
| OG0002179 | 12 | 1 | 2 | 2 | 0 | 1 | 1 | 0 | 1 | 1 | 2 | 1 | 0.001 |
| OG0008479 | 12 | 0 | 0 | 0 | 0 | 0 | 0 | 0 | 0 | 0 | 1 | 0 | 0 |
| OG0001457 | 11 | 2 | 0 | 2 | 1 | 1 | 2 | 0 | 2 | 3 | 3 | 1 | 0.001 |

| | | | | | | | | | | | | | |
|---|---|---|---|---|---|---|---|---|---|---|---|---|---|
| OG0009637 | 11 | 0 | 0 | 0 | 0 | 0 | 0 | 0 | 0 | 0 | 0 | 1 | 0.001 |
| OG0010347 | 11 | 0 | 0 | 0 | 0 | 0 | 0 | 0 | 0 | 0 | 0 | 0 | 0.002 |
| OG0010357 | 11 | 0 | 0 | 0 | 0 | 0 | 0 | 0 | 0 | 0 | 0 | 0 | 0.002 |
| OG0000179 | 10 | 9 | 3 | 4 | 7 | 8 | 7 | 1 | 9 | 9 | 2 | 3 | 0.003 |
| OG0000611 | 10 | 0 | 4 | 1 | 1 | 4 | 3 | 4 | 3 | 8 | 1 | 2 | 0.003 |
| OG0001182 | 10 | 0 | 4 | 0 | 0 | 7 | 0 | 2 | 0 | 6 | 2 | 0 | 0 |
| OG0001664 | 10 | 0 | 0 | 0 | 0 | 0 | 0 | 7 | 6 | 0 | 2 | 2 | 0.006 |
| OG0002738 | 10 | 1 | 1 | 1 | 1 | 2 | 1 | 0 | 1 | 2 | 0 | 1 | 0.006 |
| OG0010797 | 10 | 0 | 0 | 0 | 0 | 0 | 0 | 0 | 0 | 0 | 0 | 0 | 0.002 |
| OG0010805 | 10 | 0 | 0 | 0 | 0 | 0 | 0 | 0 | 0 | 0 | 0 | 0 | 0.002 |
| OG0010813 | 10 | 0 | 0 | 0 | 0 | 0 | 0 | 0 | 0 | 0 | 0 | 0 | 0.002 |
| OG0000465 | 9 | 0 | 0 | 1 | 0 | 8 | 8 | 9 | 0 | 9 | 2 | 0 | 0 |
| OG0000735 | 9 | 0 | 0 | 6 | 9 | 2 | 7 | 0 | 1 | 3 | 0 | 1 | 0 |
| OG0001358 | 9 | 1 | 0 | 1 | 1 | 2 | 4 | 0 | 3 | 4 | 3 | 1 | 0.001 |
| OG0002194 | 9 | 0 | 1 | 1 | 1 | 3 | 2 | 3 | 0 | 3 | 1 | 0 | 0.01 |
| OG0005384 | 9 | 0 | 2 | 0 | 0 | 0 | 0 | 0 | 1 | 0 | 0 | 4 | 0 |
| OG0010792 | 9 | 0 | 0 | 1 | 0 | 0 | 0 | 0 | 0 | 0 | 0 | 0 | 0.01 |

| | | | | | | | | | | | | | |
|---|---|---|---|---|---|---|---|---|---|---|---|---|---|
| OG0010804 | 9 | 0 | 0 | 0 | 0 | 0 | 0 | 0 | 0 | 0 | 1 | 0 | 0.005 |
| OG0011154 | 9 | 0 | 0 | 0 | 0 | 0 | 0 | 0 | 0 | 0 | 0 | 0 | 0.009 |
| OG0001152 | 8 | 4 | 3 | 3 | 2 | 3 | 4 | 1 | 0 | 3 | 0 | 0 | 0.004 |
| OG0003619 | 8 | 0 | 0 | 1 | 0 | 0 | 2 | 2 | 0 | 6 | 0 | 0 | 0.001 |
| OG0008481 | 8 | 0 | 1 | 0 | 0 | 0 | 0 | 0 | 0 | 3 | 0 | 1 | 0.005 |
| OG0001184 | 7 | 0 | 1 | 2 | 2 | 2 | 2 | 0 | 5 | 2 | 2 | 6 | 0.006 |
| OG0001998 | 6 | 0 | 2 | 0 | 3 | 5 | 2 | 0 | 2 | 4 | 0 | 1 | 0.001 |
| OG0002126 | 6 | 5 | 2 | 4 | 3 | 2 | 1 | 0 | 0 | 1 | 0 | 0 | 0.01 |
| Underrepresented orthogroups | *Kingdonia* | *Arabidopsis* | *Macleaya* | *Nelumbo* | *Oryza* | *Populus* | *Vitvis* | *Circaeaster* | *Akebia* | *Aqucilegia* | *Euptelea* | *Nandina* | *P* value |
| OG0000026 | 8 | 12 | 11 | 39 | 33 | 19 | 22 | 9 | 22 | 18 | 16 | 13 | 0.001 |
| OG0000048 | 5 | 24 | 14 | 7 | 13 | 10 | 8 | 11 | 8 | 29 | 10 | 6 | 0 |
| OG0000158 | 2 | 5 | 5 | 7 | 6 | 17 | 5 | 3 | 5 | 3 | 10 | 8 | 0.01 |
| OG0000099 | 1 | 15 | 7 | 21 | 27 | 8 | 7 | 3 | 3 | 1 | 4 | 4 | 0 |
| OG0000109 | 1 | 2 | 10 | 7 | 21 | 9 | 13 | 2 | 2 | 12 | 10 | 6 | 0 |
| OG0000116 | 1 | 21 | 6 | 7 | 7 | 9 | 8 | 8 | 4 | 7 | 9 | 1 | 0 |
| OG0000162 | 1 | 12 | 15 | 8 | 1 | 3 | 8 | 5 | 3 | 8 | 2 | 9 | 0 |

| | | | | | | | | | | | | | |
|---|---|---|---|---|---|---|---|---|---|---|---|---|---|
| OG0000249 | 1 | 12 | 3 | 3 | 6 | 7 | 9 | 3 | 4 | 10 | 2 | 1 | 0.004 |
| OG0000349 | 1 | 17 | 5 | 2 | 2 | 2 | 2 | 5 | 5 | 4 | 3 | 4 | 0.001 |
| OG0000390 | 1 | 11 | 1 | 11 | 6 | 1 | 2 | 2 | 6 | 2 | 3 | 3 | 0.009 |
| OG0000931 | 1 | 1 | 1 | 4 | 8 | 1 | 1 | 1 | 1 | 13 | 1 | 1 | 0.001 |
| OG0000110 | 0 | 21 | 5 | 7 | 7 | 3 | 36 | 0 | 0 | 12 | 3 | 0 | 0 |
| OG0000252 | 0 | 1 | 3 | 1 | 1 | 10 | 30 | 0 | 2 | 0 | 9 | 4 | 0 |
| OG0000267 | 0 | 13 | 0 | 4 | 11 | 2 | 12 | 0 | 2 | 2 | 7 | 6 | 0 |
| OG0000361 | 0 | 1 | 4 | 7 | 7 | 2 | 15 | 0 | 5 | 2 | 1 | 7 | 0.001 |
| OG0000440 | 0 | 2 | 18 | 0 | 0 | 13 | 3 | 4 | 0 | 0 | 3 | 4 | 0 |
| OG0000443 | 0 | 0 | 0 | 1 | 0 | 3 | 3 | 28 | 3 | 0 | 8 | 1 | 0 |
| OG0000503 | 0 | 8 | 7 | 3 | 10 | 0 | 1 | 0 | 1 | 12 | 2 | 0 | 0 |
| OG0000507 | 0 | 8 | 5 | 4 | 0 | 9 | 7 | 1 | 0 | 6 | 2 | 2 | 0.01 |
| OG0000573 | 0 | 0 | 11 | 2 | 1 | 4 | 5 | 0 | 2 | 10 | 3 | 4 | 0 |
| OG0000681 | 0 | 1 | 6 | 2 | 3 | 4 | 2 | 0 | 3 | 13 | 1 | 4 | 0.001 |
| OG0001187 | 0 | 11 | 0 | 3 | 1 | 2 | 1 | 0 | 2 | 3 | 7 | 0 | 0.001 |

Table S9. Length comparasion of *ndh* genes between *K. uniflora* and *C. agrestis.*

| | *C. agrestis* (bp) | *K. uniflora* (bp) |
|---|---|---|
| *ndhA* | 1089 | 553 |
| *ndhB* | 1530 | 723 |
| *ndhC* | 360 | 0 |
| *ndhD* | 1500 | 15 |
| *ndhE* | 303 | 303 |
| *ndhF* | 2205 | 0 |
| *ndhG* | 528 | 0 |
| *ndhH* | 1179 | 615 |
| *ndhI* | 540 | 0 |
| *ndhJ* | 474 | 468 |
| *ndhK* | 666 | 234 |

Table S10. Information of species used for phylogenetic analyses. Colored characters showing the expanded taxa in Figure S5 compared with that in Figure 3.

| Species | Family | Source |
|---|---|---|
| *Oryza sativa* L. | Poaceae | NCBI |
| *Vitis vinifera* L. | Vitaceae | NCBI |

| | | |
|---|---|---|
| *Populus trichocarpa* Torr. and Gray | Salicaceae | NCBI |
| *Arabidopsis thaliana* L. | Brassicaceae | NCBI |
| *Nelumbo nucifera* Gaertner | Nelumbonaceae | NCBI |
| *Euptelea pleiosperma* J. D. Hooker and Thomson | Eupteleaceae | 1 kp |
| *Argemone mexicana* L. | Papaveraceae | 1 kp |
| *Papaver bracteatum* Lindl. | Papaveraceae | 1 kp |
| *Capnoides sempervirens* (L.) Borkh. | Papaveraceae | 1 kp |
| *Macleaya cordata* (Willd.) R. Br. | Papaveraceae | NCBI |
| *Akebia trifoliata* (Thunberg) Koidzumi | Lardizabalaceae | 1 kp |
| *Circaeaster agrestis* Maxim. | Circaeasteraceae | Current study |
| *Kingdonia uniflora* Balf. f. and W.W. Sm. | Circaeasteraceae | Current study |
| *Hydrastis canadensis* L. | Ranunculaceae | 1 kp |
| *Aquilegia coerulea* E. James | Ranunculaceae | NCBI |
| *Podophyllum peltatum* L. | Berberidaceae | 1 kp |
| *Nandina domestica* Thunberg | Berberidaceae | 1 kp |

**Figure S1.** Distribution range of *K. uniflora.* The black dot shows the individuals with previous occurrence record but no longer have extant populations; the red dots represent the current distribution range of *K. uniflora*.

**Figure S2.** Morphological features of *K. uniflora.*

**Figure S3**. Estimation of *K. uniflora* genome size based on flow cytometer analysis. The above panel showing the 2C DNA of *Actinidia chinensis* (Hopping, 1994) at 32377.78, and the panel below indicating 2C DNA of *K. uniflora* at 47614.17.

**Figure S4.** 17 *k*-mer frequency distribution of sequencing reads.

**Figure S5.** Comparative analyses of genomic features between *Kingdonia uniflora* and *Arabidopsis thaliana*. Tracks from inside to outside are collinearity between both genomes, number of chromosomes/scaffolds, gene density, GC content and TE density.

**Figure S6**. Dated phylogeny for 17 plant species with *Oryza* as an outgroup. A time scale is shown at the bottom.

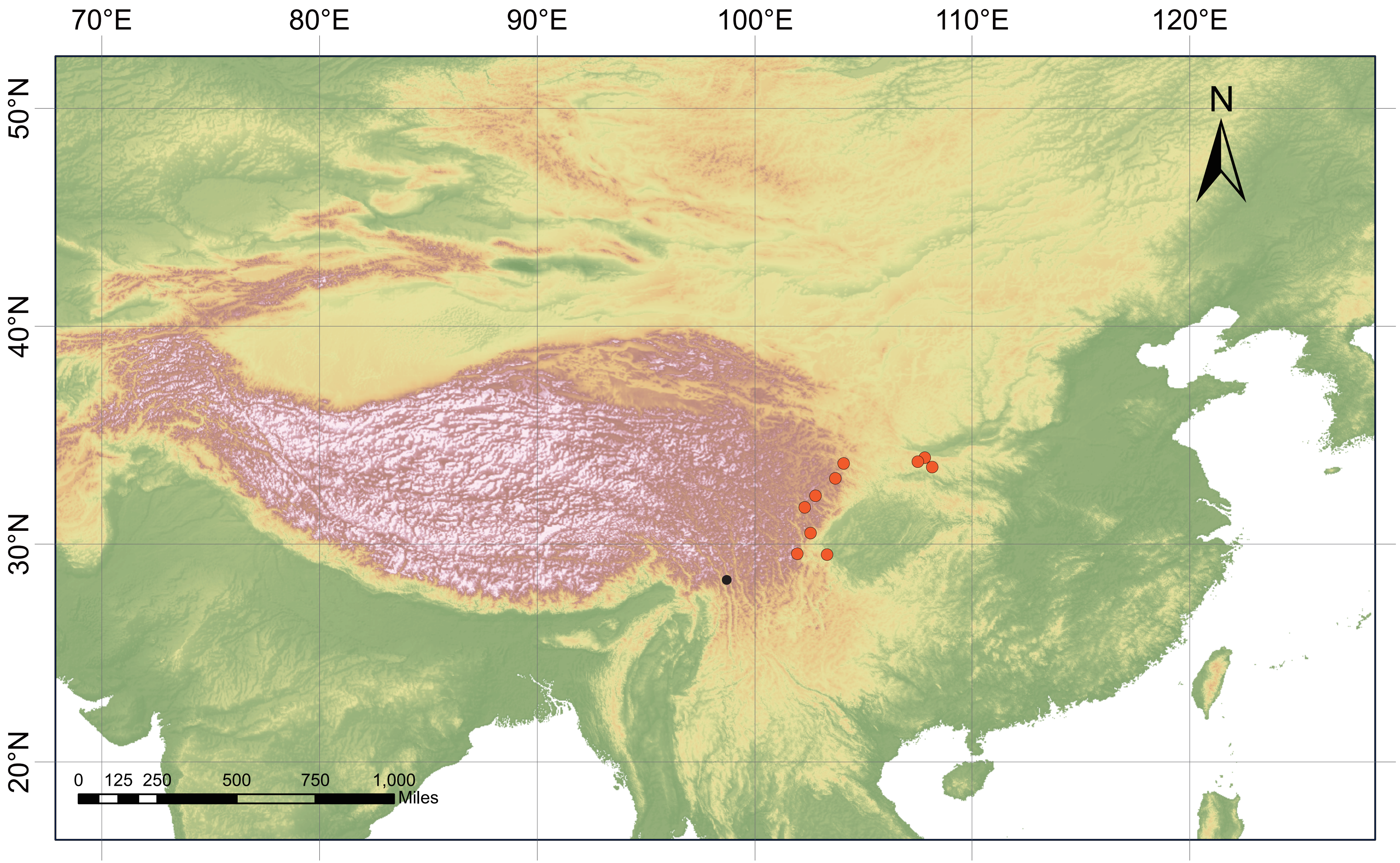
70°E
80°E
90°E
100°E
110°E
120°E
50°N
40°N
30°N
20°N
N
0 125 250 500 750 1,000
Miles

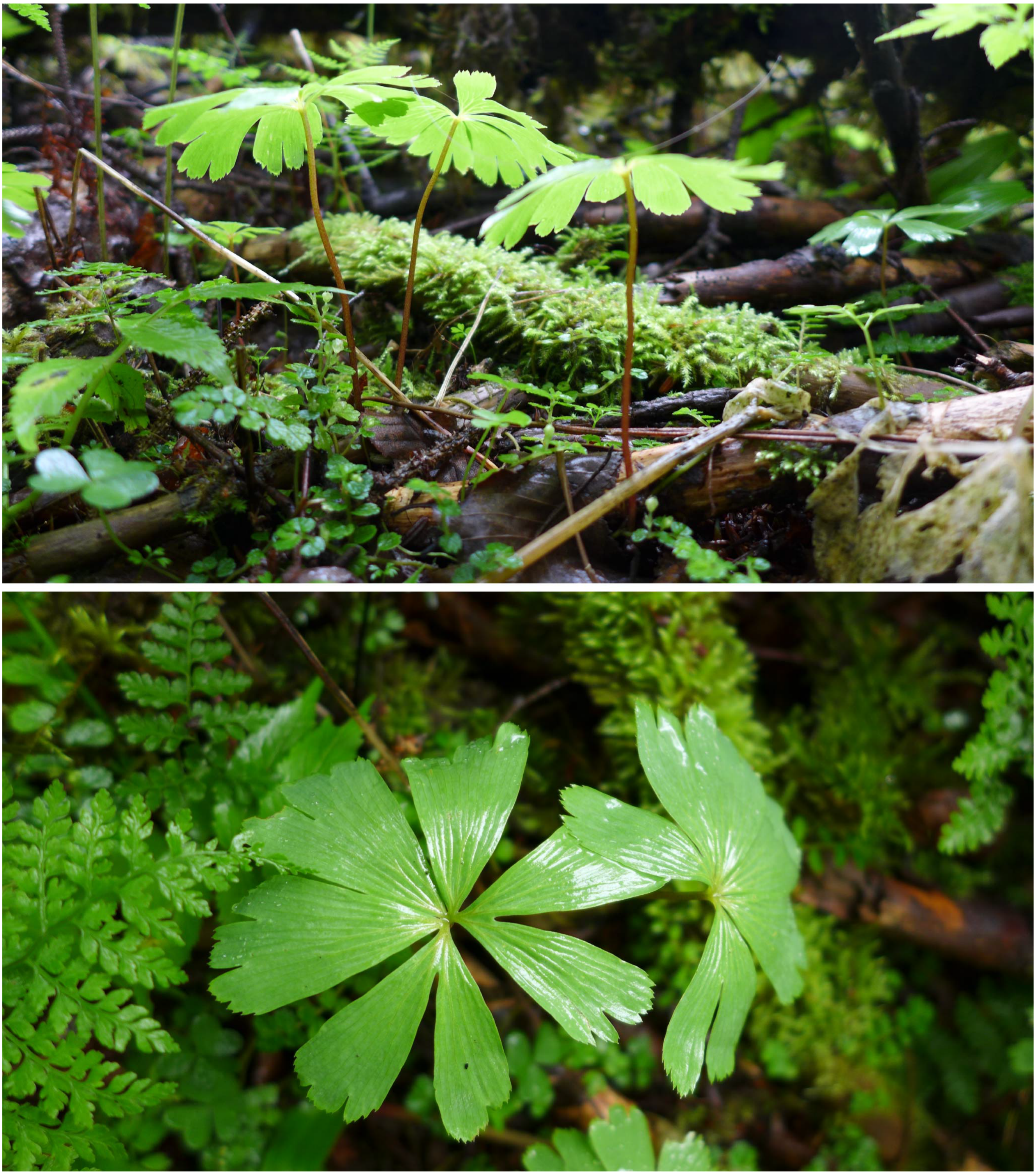

Figure S2

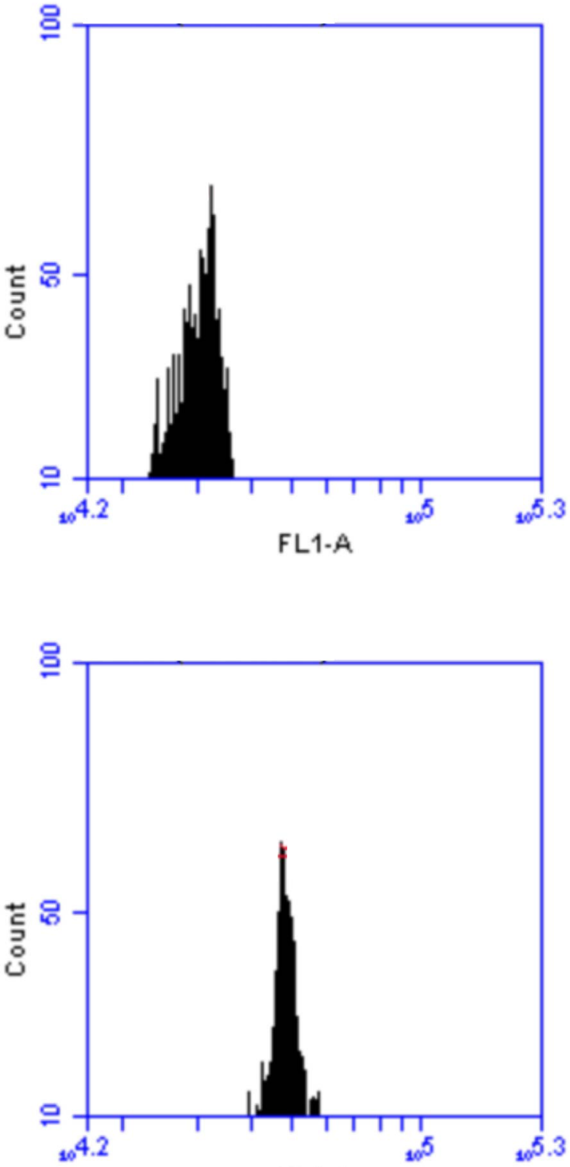
100
50
10
Count
10^4.2
10^5
10^5.3
FL1-A
100
50
10
Count
10^4.2
10^5
10^5.3
FL1-A

Figure S3

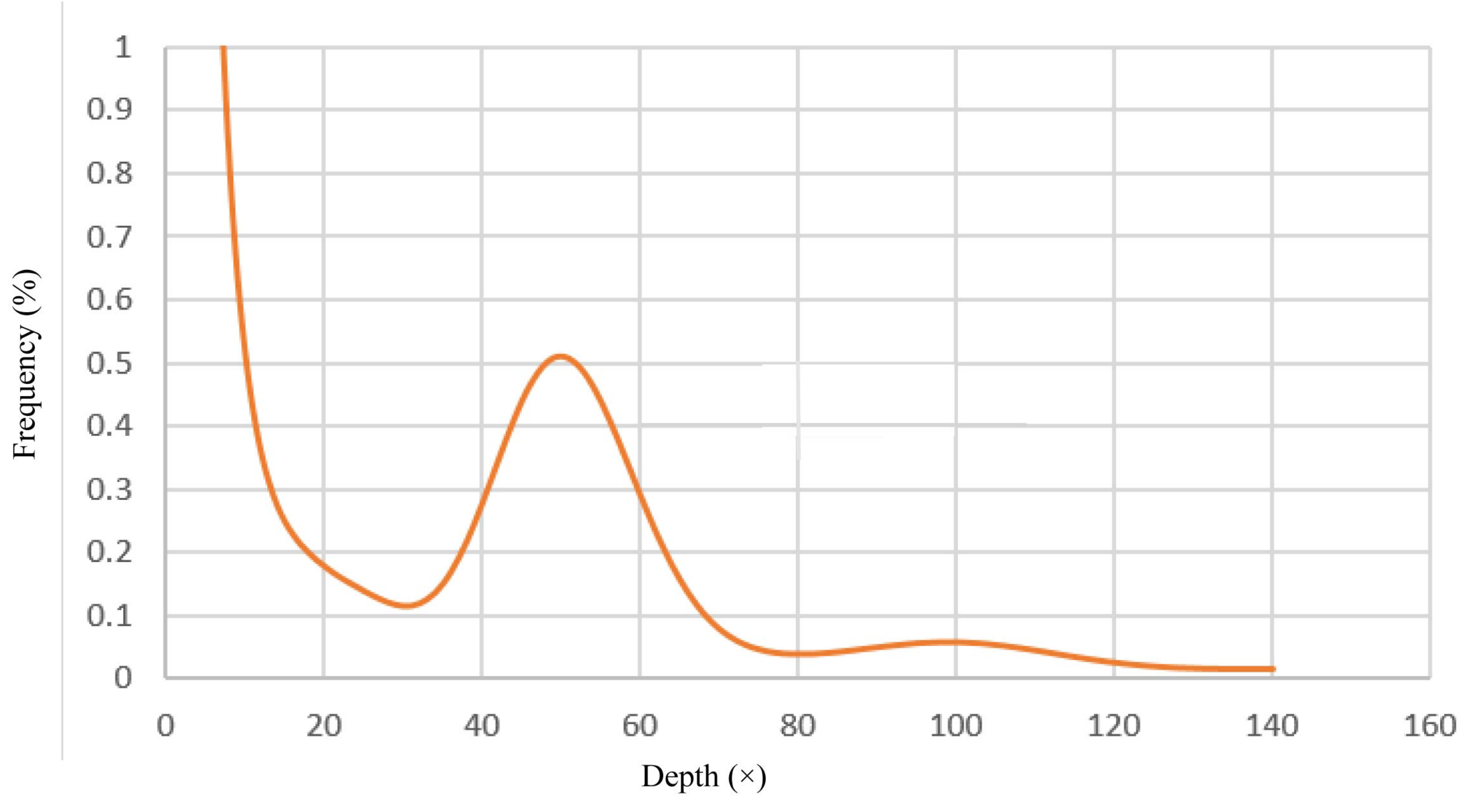
Frequency (%)
1
0.9
0.8
0.7
0.6
0.5
0.4
0.3
0.2
0.1
0
0
20
40
60
80
100
120
140
160
Depth (×)

Figure S4

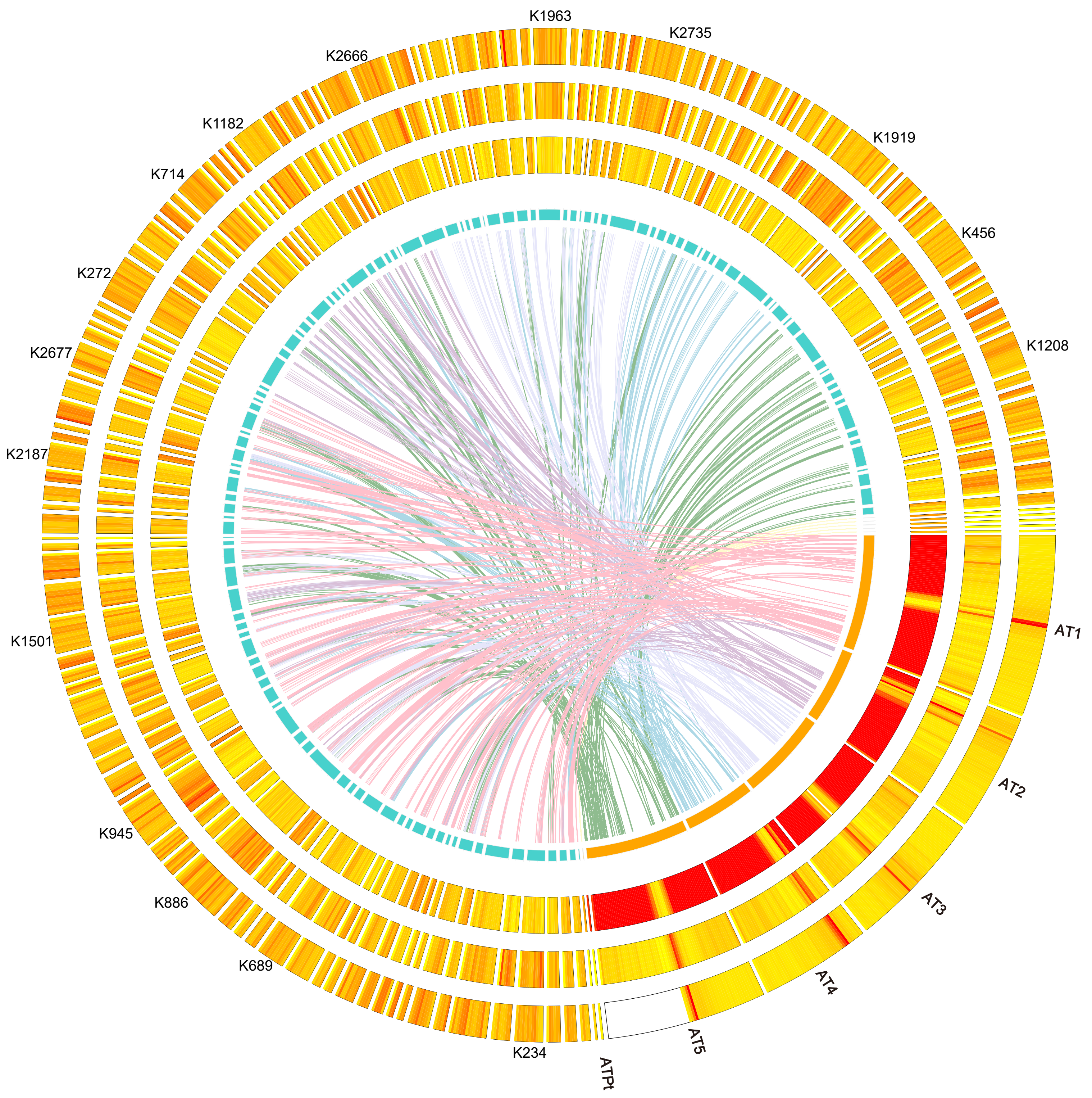
K1963
K2735
K2666
K1919
K1182
K456
K714
K1208
K272
K2677
K2187
AT1
K1501
AT2
K945
AT3
K886
AT4
K689
AT5
K234
ATPt

Figure S5

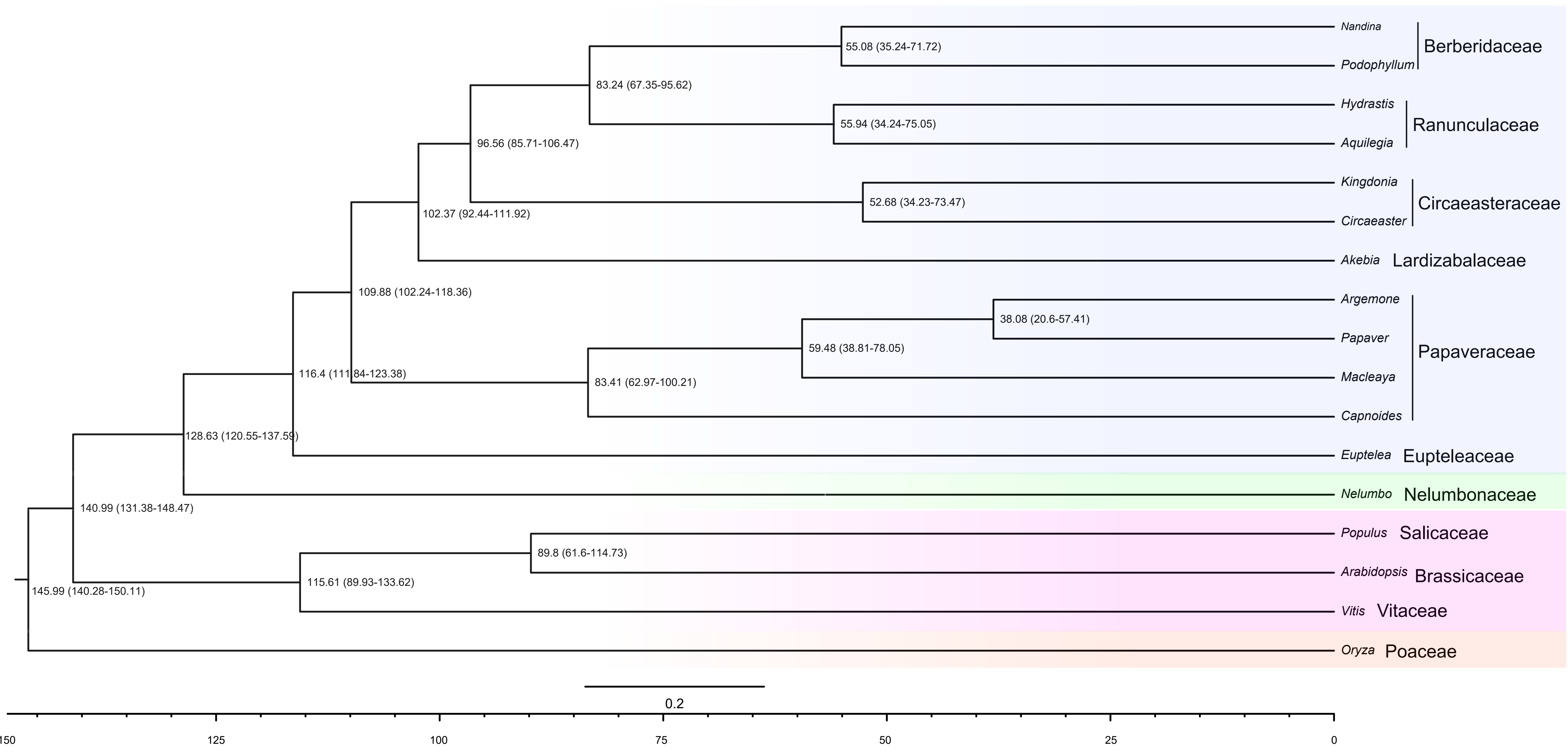

Nandina
Podophyllum
Berberidaceae
Hydrastis
Aquilegia
Ranunculaceae
Kingdonia
Circaeaster
Circaeasteraceae
Akebia
Lardizabalaceae
Argemone
Papaver
Macleaya
Capnoides
Papaveraceae
Euptelea
Eupteleaceae
Nelumbo
Nelumbonaceae
Populus
Salicaceae
Arabidopsis
Brassicaceae
Vitis
Vitaceae
Oryza
Poaceae
55.08 (35.24-71.72)
83.24 (67.35-95.62)
55.94 (34.24-75.05)
96.56 (85.71-106.47)
52.68 (34.23-73.47)
102.37 (92.44-111.92)
109.88 (102.24-118.36)
38.08 (20.6-57.41)
59.48 (38.81-78.05)
116.4 (111.84-123.38)
83.41 (62.97-100.21)
128.63 (120.55-137.59)
140.99 (131.38-148.47)
89.8 (61.6-114.73)
115.61 (89.93-133.62)
145.99 (140.28-150.11)
0.2
150
125
100
75
50
25
0


Figure S6
